# Phenotype-Guided In Silico Molecular Generation Using Large Language Models

**DOI:** 10.64898/2026.01.03.697483

**Authors:** Qun Jiang, Xing Ye, Zekun Guo, Yingce Xia, Zequn Liu, Jiaying Xu, Peiran Jin, Fusong Ju, Huanhuan Xia, Shangya Feng, Rui Jiang, Haiguang Liu, Tao Qin, Pan Deng, Sida Shao

## Abstract

Complex diseases often emerge from coordinated, system-level cellular state changes that are difficult to address with target centric drug discovery. Phenotypic drug discovery offers a principled alternative but remains constrained by the cost and scalability of pathologically relevant assays. Here we present GEMGen, a large language model–based framework that performs in silico phenotypic drug discovery by generating small molecules directly from transcriptomic representations of cellular states. GEMGen encodes desired phenotypic transitions as text-based representations of up- and down-regulated gene sets, enabling transferable modeling across experimental platforms and data modalities. Trained on large scale chemical perturbation data, GEMGen robustly identifies phenotype-oriented compounds and mechanistically related but structurally distinct candidates across multiple benchmarks. Applied to signatures induced by genetic perturbations, GEMGen produces small molecules that phenocopy gene knockdown effects and identifies chemically novel inhibitors, including previously unreported KEAP1 inhibitors that activate NRF2 signaling. Extending this approach to a disease relevant model of fibrosis, GEMGen generates compounds that reverse profibrotic transcriptional programs and cellular phenotypes. These results establish a scalable framework for translating transcriptomic phenotypes into candidate therapeutic molecules, enabling systems-level exploration of vast chemical space and offering a complementary *in silico* counterpart to physical phenotypic drug screens.

## Introduction

Disease phenotypes often arise from coordinated, system level alterations across gene regulatory programs, signaling networks, and cellular state transitions rather than from isolated molecular lesions (1). As a result, direct causal links between individual molecular targets and observed phenotypic abnormalities are frequently obscured. This complexity poses a fundamental challenge for therapeutic discovery, particularly when pathogenic mechanisms are multifactorial or incompletely understood (2,3).

Phenotypic drug discovery (PDD) addresses this challenge by prioritizing compounds based on their ability to induce desired cellular or tissue level outcomes, independent of predefined etiological targets, offering an alternative entry point for therapeutic development (4–6). Historically, this strategy has yielded a disproportionate number of first-in-class medicines, especially in disease contexts characterized by complex, emergent phenotypes that are poorly captured by reductionist, single target approaches (7–10). However, classical phenotypic screen relies on complex, disease relevant experimental models that are costly, time consuming, and difficult to standardize. These constraints are further amplified by the need to screen large chemical libraries, ultimately limiting the scalability and accessibility of traditional PDD approaches (5,6).

In recent years, advancement in high-throughput transcriptomic profiling have established gene expression as a practical and information rich proxy for cellular phenotype, enabling phenotypic states to be measured at scale. In parallel, progress in machine learning has motivated the development of computational approaches to perform phenotypic screen *in silico* using transcriptomic data. For example, multiple methods have been proposed to rank or screen compounds (11–16), or to design new molecules (17–21), based on their ability to reverse disease associated transcriptional signatures derived from perturbational datasets. Despite these advances, transcriptome driven drug discovery remains challenged by technical heterogeneity across platforms, batch effects, and variable data quality, which often limit model generalization across studies and experimental systems. Furthermore, the *de novo* generation of chemically valid structures that exhibit high drug-likeness is inherently complex. As a result, relatively few computational methods have produced compounds that translate robustly to wet lab validation, underscoring the need for transferable frameworks that can propose therapeutically relevant molecules with new chemistry and real-world applicability.

To address these limitations, we developed GEMGen (Gene Expression–guided Molecular Generator), a large language model–based framework for *in silico* phenotypic drug generation. Rather than operating on raw quantitative expression profiles, GEMGen represents transcriptomic perturbations as ranked lists of up- and down- regulated genes derived from differential expression analyses between disease and control states. This abstraction preserves the directionality and relative importance of transcriptional changes while mitigating platform specific biases, enabling improved transferability across experimental modalities with minimal loss of biological context (22–24). GEMGen then conditions *de novo* molecular generation on these textual representations, allowing exploration of chemical space beyond existing libraries and prior target annotations (25–27).

We trained GEMGen on the LINCS L1000 chemical perturbation dataset, which profiles transcriptional responses to thousands of small molecules using a scalable bead-based assay (4). When evaluated on held out compounds, GEMGen generated chemically distinct molecules not observed during training, demonstrating generalization beyond memorized structures. We further assessed cross platform robustness using the Tahoe 100M single cell chemical perturbation dataset (20), showing that GEMGen can operate effectively across bulk and single cell transcriptomic modalities. Applied to genetic perturbation settings (21), GEMGen recovered known inhibitors of the perturbed genes while also proposing previously uncharacterized molecules that recapitulated the expected transcriptional shifts in experimental assays, revealing alternative strategies for modulating gene driven phenotypes. Finally, applying GEMGen to a disease relevant model of fibrosis, we identified compounds that reversed profibrotic transcriptional programs and attenuated extracellular matrix production, demonstrating the ability of GEMGen to translate complex pathological gene expression signatures into functional therapeutic candidates.

Together, these results establish GEMGen as a generalizable framework for transcriptome guided phenotypic drug generation, bridging large scale perturbational data and disease relevant phenotypes. By complementing experimental disease models with *in silico* generative design, GEMGen enables scalable, systems level therapeutic discovery for complex diseases.

### Overview of GEMGen

GEMGen is a generative large language model (LLM) framework that proposes candidate compounds from transcriptomic representations of cellular states (**Fig. 1a**). Given paired transcriptomes describing healthy and diseased cells, GEMGen generates small molecules predicted to induce the desired phenotypic transition (**Fig. 1b**).

**Figure 1.**
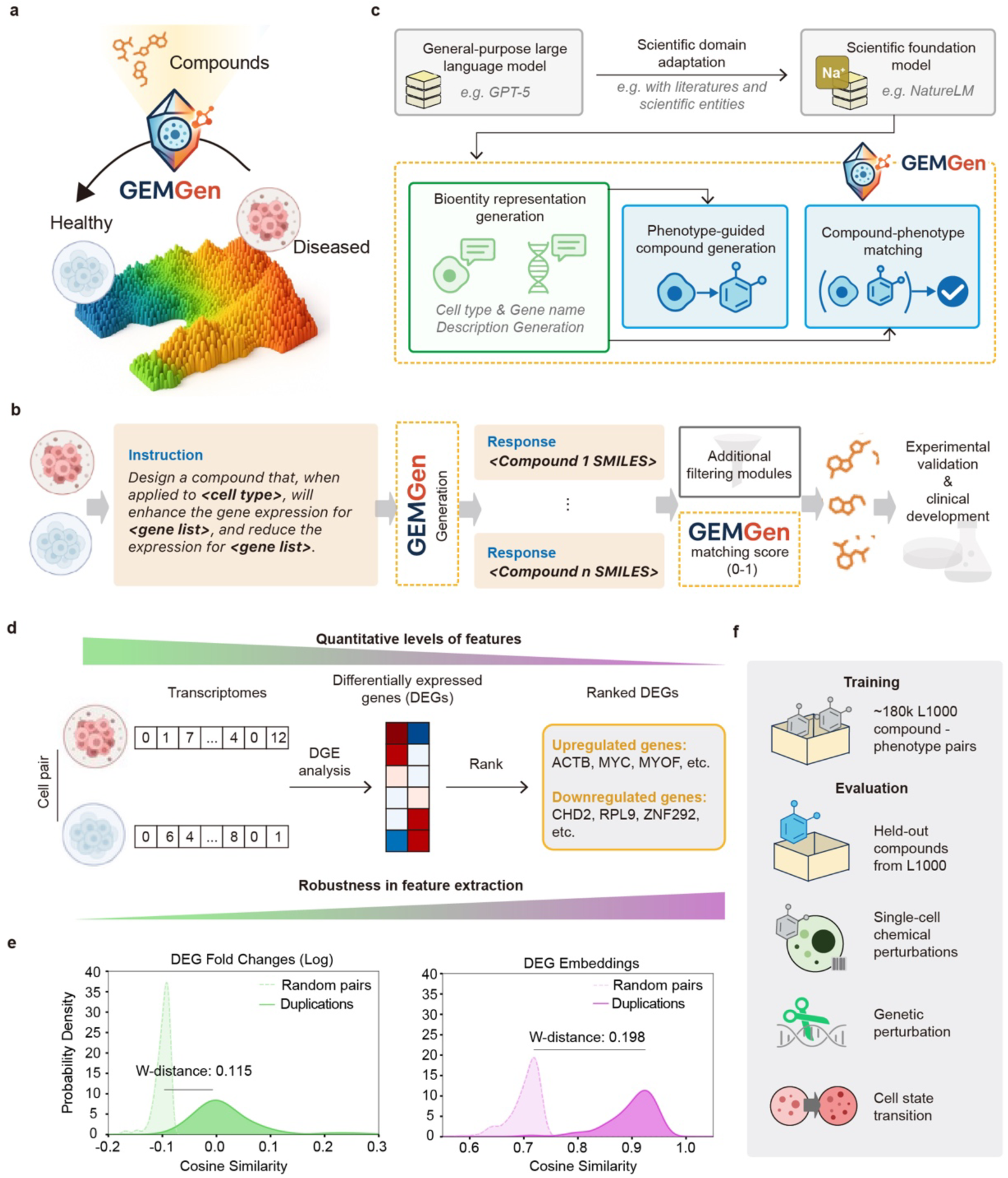
Overview of the GEMGen framework. a, Conceptual illustration of GEMGen. Given transcriptomic profiles describing transitions between healthy and diseased cellular states, GEMGen generates candidate compounds predicted to induce the desired phenotypic change. **b,** End-to-end workflow of GEMGen. Transcriptomic changes are provided as text-based, rank-ordered lists of up- and down-regulated genes within a specific cellular context. GEMGen generates candidate compounds in SMILES format, which are optionally filtered by additional modules and ranked using the GEMGen scorer to prioritize candidates for experimental validation and downstream development. **c**, Model architecture and training strategy. GEMGen builds on a scientific foundation model (e.g., NatureLM) adapted from a general-purpose large language model (e.g., GPT-5). On this basis, GEMGen performs bioentity representation generation, followed by phenotype-guided compound generation and compound–phenotype matching. **d**, Transformation of quantitative transcriptomic measurements into standardized, text-based features. Paired transcriptomes are processed through differential gene expression (DGE) analysis and converted into ranked lists of up- and down-regulated genes. **e,** Rank-based representations improve robustness while preserving phenotypic signal. Distributions of cosine similarity for numeric DEG fold changes (left) and text-based DEG embeddings (right) are shown for matched treatments and random pairs across the L1000 and Tahoe-100M datasets. Rank-based representations yield greater separation between matched and random pairs, as quantified by the Wasserstein distance (W-distance), indicating improved robustness for downstream modeling. **f**, Training and evaluation settings. GEMGen is trained on approximately 180,000 compound–phenotype pairs from the L1000 dataset and evaluated across multiple scenarios, including held-out compounds, single-cell chemical perturbations, genetic perturbations, and disease-associated cell state transitions.

GEMGen builds upon a science-focused LLM trained on a curated corpus spanning natural language, scientific literature, and structured biological databases (28) (**Fig. 1c**). This scientific foundation model operates in an instruction–response format, enabling the interpretation of text-based biological prompts and the generation of contextually relevant outputs across biological modalities. GEMGen extends this foundation to enable phenotype-driven drug discovery through three complementary instruction–response tasks.

The first, a bioentity representation task, trained the model to generate textual descriptions of genes and cell types curated from the NCBI database and the ATCC cell catalog (**Fig. 1c and Fig. S1, Methods**). Joint training on gene- and cell-type-level samples encouraged a coherent semantic representation space, thereby facilitating biologically meaningful interpretation of transcriptomic changes. This model served as the basis for subsequent tasks.

The second task, phenotype-guided compound generation, trained the model to generate chemical structures capable of eliciting a given transcriptomic change (**Fig. 1c**). The model received ranked lists of genes up- and down-regulated representing the desired phenotypic transition and was tasked with generating compounds predicted to induce matching transcriptomic responses. For training, we used the large-scale LINCS L1000 dataset, which provides gene expression signatures for over one million perturbation experiments spanning approximately 30,000 small molecules across diverse human cell types (12) (**Methods**). These data establish broad correspondences between chemical structures and their induced cellular transcriptional states for GEMGen.

In parallel, a compound–phenotype matching task trained a scoring model to estimate the likelihood that a given compound reproduces a target gene expression signature. This scorer outputs a continuous match score between 0 (no match) and 1 (perfect match), enabling prioritization of generated compounds for experimental validation (**Fig. 1b–c**). The scorer was trained independently using the same L1000 dataset, with negative samples generated by pairing compounds with unrelated expression profiles (21) **(Methods)**.

For the latter two tasks, quantitative transcriptomic measurements were transformed into text-based ranked lists of differentially expressed genes (**Fig. 1d**), providing a standardized textual abstraction of cellular states. This representation aligns naturally with the token-based operation of LLMs. More importantly, text-based descriptions abstract away dataset-specific expression magnitudes while preserving relative transcriptional changes, resulting in greater consistency across transcriptomic datasets and improved robustness of phenotypic representations. This is exemplified by greater separation based on Wasserstein distance between randomized pairs and matched treatments using text-based representations than with numeric differential expression values (**Fig. 1e, Methods**). Consequently, although trained on L1000 bulk expression profiles, GEMGen holds promise to generalize effectively to transcriptomic data generated across diverse settings and application scenarios (**Fig. 1f**).

Together, these three tasks equip GEMGen with complementary capabilities in bioentity representation, phenotype-guided molecular generation, and compound prioritization, forming a unified generative framework for in silico phenotypic drug discovery.

### Benchmarking GEMGen for phenotype-guided molecular generation

To benchmark GEMGen’s phenotype-guided molecular generation, we assembled two complementary evaluation datasets, each probing a distinct aspect of model performance. First, we held out 404 L1000 samples as a “novel compound” test set, in which each compound has a maximal Tanimoto similarity below 0.6 based on Morgan fingerprint to any training molecules (**Fig. 2a; Methods**). This setting evaluates whether GEMGen can generate compounds from previously unseen chemical space rather than recalling known chemical structures. Second, to assess generalization across experimental conditions, we curated 640 high-quality compound–phenotype pairs from the Tahoe-100M dataset (29) (**Fig. 2a**), a large-scale single-cell perturbation atlas profiling transcriptional responses to small molecules across 50 cancer cell lines (**Methods**). Unlike bulk L1000 data, which captures population-averaged expression changes, Tahoe-100M measures heterogeneous responses at single-cell resolution. This novel condition test therefore evaluates whether a model trained exclusively on bulk transcriptomic profiles can transfer to a distinct measurement modality and experimental regime.

**Figure 2.**
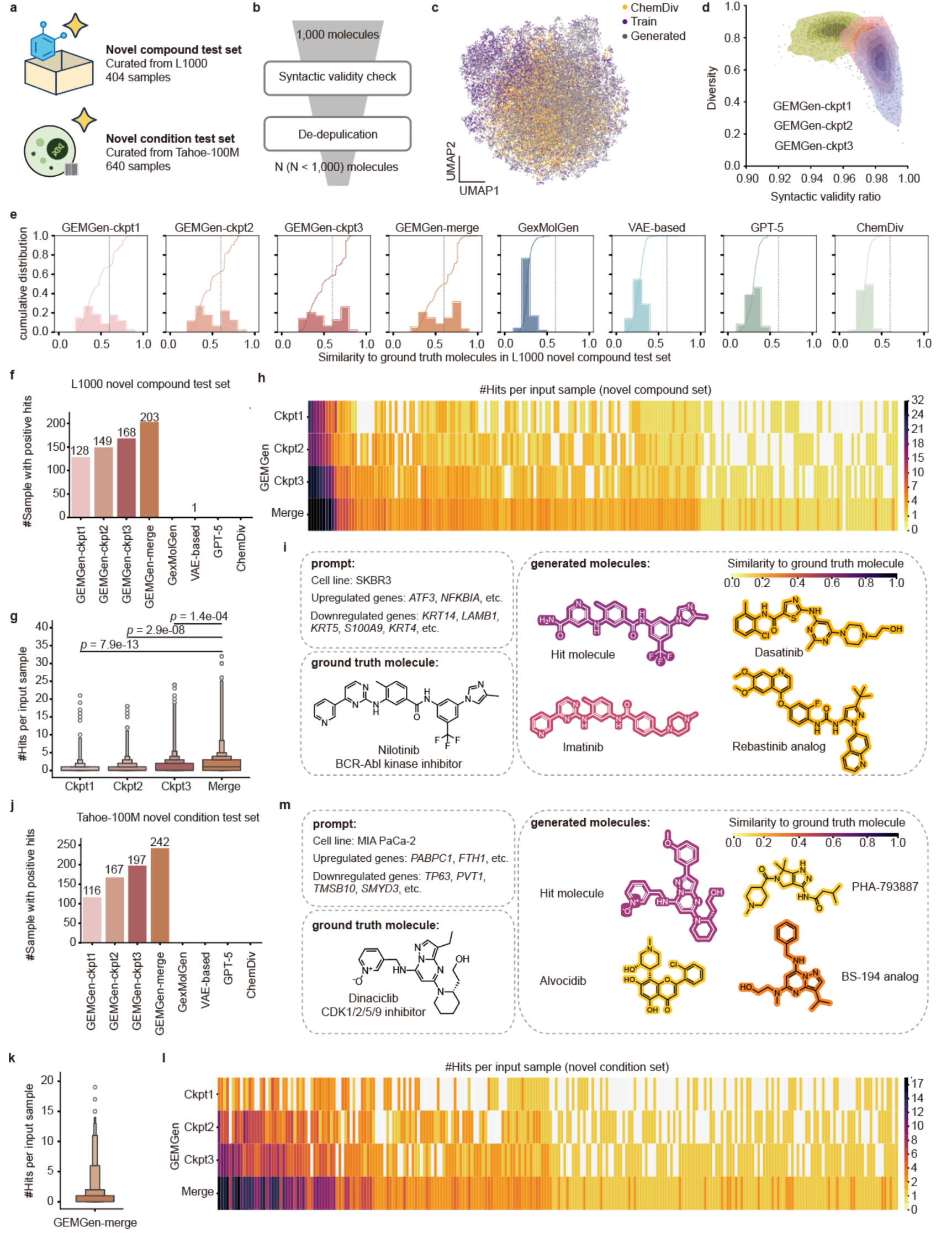
GEMGen generalizes across novel compounds and sequencing conditions. **a**, Two complementary evaluation datasets were constructed: a novel compound test set curated from L1000 (404 samples) and a novel condition test set curated from Tahoe-100M (640 samples). **b**, Workflow of compound generation and filtering. For each input phenotype, GEMGen generated 1,000 candidate molecules in SMILES format, which were filtered for syntactic validity and de-duplicated before downstream analysis. **c**, UMAP visualization of chemical space spanned by GEMGen-generated compounds, ChemDiv compounds, and training compounds. For visualization, 10,000 molecules were randomly sampled from each set. **d**, Relationship between syntactic validity and compound diversity across three GEMGen variants (Ckpt1–3) trained for increasing numbers of epochs. **e**, Cumulative distribution of Tanimoto similarity between generated and their corresponding ground-truth compounds in the L1000 novel compound test set. Dashed lines indicate a similarity threshold of 0.6. **f**, Number of phenotypic conditions with at least one positive hit in the L1000 novel compound test set under similarity thresholds of 0.6. The ensemble model (“merge”), which combines all checkpoints, yields more hits than any individual variant or baseline method. **g**, Distribution of hit counts per condition in the L1000 test set. Ensemble predictions show significantly higher numbers of hits. *P-*values are calculated with two-sides Mann-Whitney U test. *N*=404. **h**, Per-condition heatmap of hit counts in the L1000 test set. **i**, Representative example from the L1000 test set, showing input phenotype, ground-truth compound, scaffold-level similar hit, and mechanistically relevant but structurally distinct compounds. **j**, Number of phenotypic conditions with at least one positive hit in the Tahoe-100M novel condition test set under similarity thresholds of 0.6. **k**, Distribution of hit counts per condition in the Tahoe-100M test set. *N*=640. **l**, Per-condition heatmap of hit counts in the Tahoe-100M test set. **m**, Representative example from the Tahoe-100M test set, showing input phenotype, ground-truth compound, scaffold-level similar hit, and mechanistically relevant but structurally distinct compounds.

For each input phenotype, GEMGen generated 1,000 candidate compounds in SMILES format predicted to reproduce the corresponding gene expression changes. Generated molecules were filtered for syntactic validity and de-duplicated prior to analysis (**Fig. 2b**). Compared with training compounds and ChemDiv, a large-scale chemical library comprising over 1.5 million real molecules, GEMGen-generated molecules span both close analogs of known compounds and structurally distinct chemotypes (**Fig. 2c and Fig. S2**), indicating effective exploration of chemical space beyond the training distribution.

We trained three GEMGen variants with increasing numbers of training epochs. As expected, longer training improved syntactic validity while reducing molecular diversity (**Fig. 2d and Fig. S3a–b**). All variants maintained comparable drug-likeness, assessed via Lipinski’s Rule of Five (Ro5) (30), quantitative estimate of drug-likeness (QED) (31), and synthetic accessibility score (SAS) (32) (**Fig. S3c-e**). We therefore retained all three models for downstream analyses and combined their outputs into an ensemble pool (GEMGen-merge).

We next evaluated whether GEMGen could recover compounds structurally related to ground-truth molecules associated with each phenotype. Generated molecules spanned a broad range of Tanimoto similarity values, with most falling between 0.2 and 0.8 relative to the corresponding ground-truth structures (**Fig. 2e**). Using a similarity threshold of 0.6 to define positive hits, each individual GEMGen variant recovered at least one ground-truth–related compound in over 100 phenotypic conditions (**Fig. 2f**), whereas the ensemble model (GEMGen-merge) achieved such recovery in more than 200 conditions (**Fig. 2e–f**). In addition, GEMGen-merge yielded a higher number of hits per condition compared with any single model (**Fig. 2g–h**). These results indicate that GEMGen can consistently generate compounds structurally related to known actives across diverse phenotypic contexts, and that ensemble generation further expands coverage of plausible candidate structures, thereby improving discovery efficiency.

As similarity-based thresholds capture only scaffold-level resemblance, the reported hit rates likely underestimate GEMGen’s recovery performance. In addition to close structural analogs of ground-truth compounds, GEMGen can generate molecules with low Tanimoto similarities that are known to act on the same molecular targets or pathways. For example, when conditioned on nilotinib induced transcriptomic signatures, GEMGen generated not only nilotinib analog, but also other ABL kinase inhibitors such as imatinib, dasatinib and a rebastinib analog. (**Fig. 2i**). This observation suggests that GEMGen captures phenotype–target relationships beyond scaffold-level similarity, enabling the discovery of mechanistically relevant but structurally distinct candidates.

We further compared GEMGen against four representative alternative approaches **(Fig. S4)**. We first implemented a variational autoencoder (VAE)–based phenotype-guided molecular generator, following the design principles of Gex2SGen and trained on the same data as GEMGen (**Methods**). VAE-based architectures are widely used in molecular design for their smooth latent spaces and interpolation capabilities (19,33–35). We also evaluated GexMolGen (17), which builds on the single-cell foundation model scGPT (36), and directly encodes gene-level expression profiles for molecule generation. In addition, we tested a general-domain large language model (GPT-5o) prompted with expression-derived instructions but without domain-specific fine-tuning, to assess whether generic LLMs can infer biologically meaningful molecules from transcriptional patterns alone. Finally, we included random sampling from ChemDiv as a reference baseline representing unguided exploration of chemical space. Across the novel compound test set, none of the baseline approaches generated appreciable numbers of compounds structurally related to ground-truth molecules (**Fig. 2e–f**). In contrast, all three GEMGen variants consistently recovered substantially more hits, with the ensemble model (GEMGen-merge) achieving the strongest overall performance (**Fig. 2e–h**).

This performance advantage extended to the novel condition test set derived from single-cell chemical perturbation data. When evaluated on Tahoe-100M, GEMGen again outperformed all alternative approaches by a wide margin (**Fig. 2j–l**). Notably, whereas conventional deep generative models and general-domain LLMs struggled to produce biologically relevant molecules under cross-platform distribution shift, GEMGen maintained its ability to recover compounds closely related to known actives while also generating mechanistically relevant but structurally distinct candidates (**Fig. 2m**). Together, these results indicate that GEMGen learns a robust phenotype–compound mapping that generalizes across substantial experimental condition shifts.

### Scoring-based prioritization of GEMGen-generated compounds

Experimentally testing thousands of generated molecules is impractical, particularly because many candidates are not available in commercial libraries. We therefore introduced a compound–phenotype scoring model to prioritize GEMGen-generated molecules according to their likelihood of reproducing a given transcriptional signature. This scoring step is designed to complement generation by enriching for biologically relevant candidates before experimental validation.

We first evaluated whether the scorer could discriminate true compound–phenotype pairs from randomly shuffled negatives (**Methods**). Across the combined novel compound and novel condition test sets, the scorer achieved an area under the receiver operating characteristic curve (AUROC) of 0.939 and an area under the precision–recall curve (AUPRC) of 0.945 (**Fig. 3a–b**). The score distributions showed strong separation between negative (median = 0.085) and positive (median = 0.849) pairs (**Fig. 3c**). At a score threshold of 0.8, precision, reflecting the fraction of predicted positives that correspond to true matches, reached 0.950 (**Fig. 3d**), demonstrating that the scorer provides a reliable signal for compound–phenotype matching.

**Figure 3.**
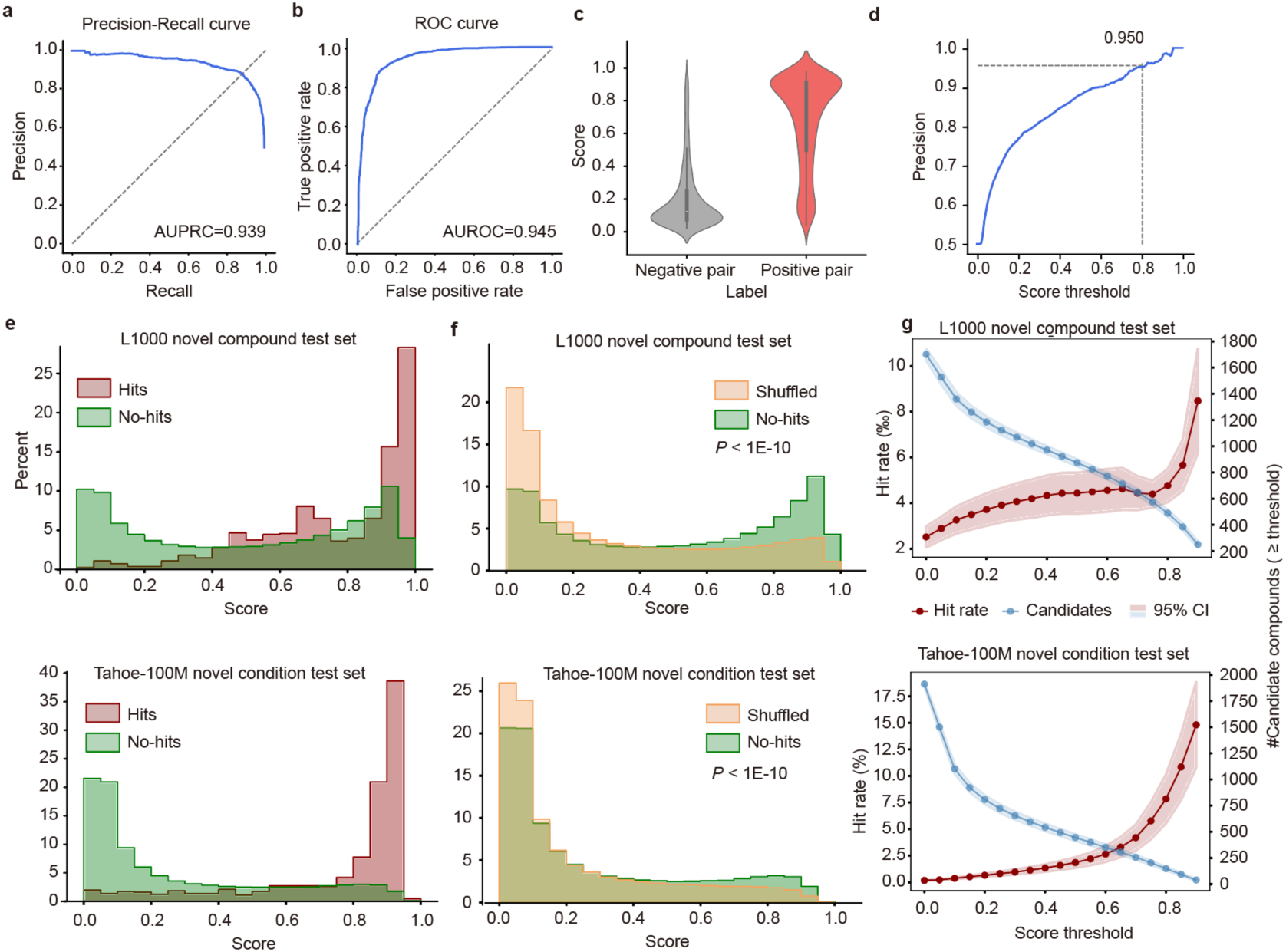
Scoring model distinguishes true compound–phenotype pairs and prioritizes effective candidates. **a–b**, Performance of the scorer in distinguishing true compound–phenotype pairs from random negatives, evaluated by (a) precision–recall and (b) receiver operating characteristic (ROC) curves. **c**, Distribution of scores assigned to positive and negative pairs. The scorer assigns significantly higher values to true pairs (two-sided Mann-Whitney U test, P < 1×10^-15^). N=2,088 for both positive and negative pairs. **d**, Precision at different score threshold. Dashed line: a precision of 0.950 is achieved at a threshold of 0.8. **e,** Score distributions for compounds generated in the L1000 (top) and Tahoe-100M (bottom) test sets. “Hits” are generated molecules with Tanimoto similarity >0.6 to ground-truth compounds; “non-hits” are dissimilar molecules. The scorer assigns higher scores to hits than to non-hits. **f,** Score distributions for generated compounds versus randomly shuffled compounds in the L1000 novel compound and Tahoe-100M novel condition test sets. *P-*values are calculated with two-sides Mann-Whitney *U* test. **g**, Effect of filtering by score threshold on candidate retention and hit rate in the L1000 (top, N=203) and Tahoe-100M (bottom, N=242) test sets. Shaded bands: 95% confidence intervals.

Applying the scorer to GEMGen-generated compounds revealed a clear correspondence between structural proximity and predicted phenotypic relevance. Compounds with Tanimoto similarity greater than 0.6 to ground-truth molecules (“hits”) received significantly higher scores than those with lower similarity (“non-hits”; **Fig. 3e**). Importantly, because ground-truth compounds are not the only valid solutions, we further examined non-hit compounds. When compared with randomly paired compound–phenotype controls (**Methods**), non-hits exhibited significantly higher match scores than random background (**Fig. 3f**), indicating that GEMGen generates chemically novel molecules that are nonetheless predicted to induce similar transcriptional responses.

We next asked whether score-based filtering improves practical compound discovery. In the novel condition test set, increasing the score threshold progressively reduced the number of candidate molecules while monotonically increasing hit rates (**Fig. 3g and Fig. S5a**). A similar trend was observed in the novel compound test set, although hit rates declined above a threshold of 0.6 (**Fig. 3g and Fig. S5b**), likely reflecting reduced scorer confidence for highly novel chemical structures. To further assess ranking performance, we performed a top-K analysis using score-ranked ensemble GEMGen outputs (∼2,000 molecules per phenotype). As K decreased, the proportion of true hits increased steadily (**Fig. S5c–d**), demonstrating that higher-scoring compounds are preferentially enriched for ground-truth matches. Together, these results establish the compound–phenotype scorer as an effective prioritization module that improves the efficiency of GEMGen-guided compound discovery.

### GEMGen generates compounds that phenocopy genetic perturbations

Most therapeutic small molecules act by inhibiting target activities, thereby reproducing loss-of-function phenotypes at the gene level. We therefore evaluated whether GEMGen can generate compounds that phenocopy genetic perturbations by recapitulating transcriptional signatures induced by targeted gene knockdown. In this setting, the model is tasked with producing small molecules whose predicted phenotypic effects resemble those of specific genetic perturbations (**Fig. 4a**).

**Figure 4.**
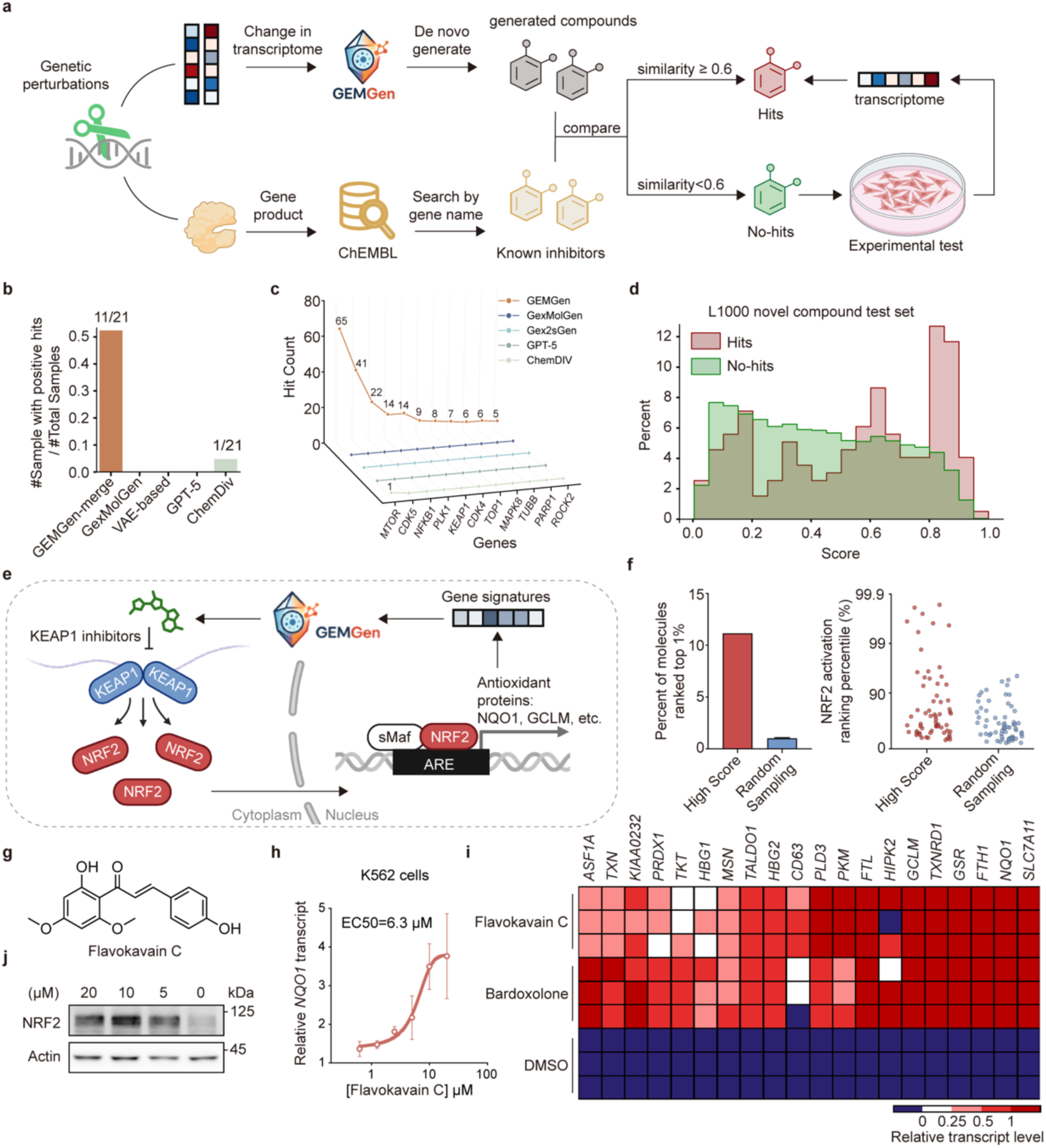
GEMGen can generate compounds directly from genetic perturbation signatures. a, Schematic overview of the evaluation framework for genetic perturbations. Genetic perturbations induce characteristic transcriptional changes that are used as input to GEMGen for de novo generation of candidate small molecules. Generated compounds are compared with known inhibitors retrieved from ChEMBL for each target. Compounds exhibiting high structural similarity to the curated ground truth molecules (similarity ≥ 0.6) are classified as hits, whereas those with lower similarity (similarity < 0.6) were designated as non-hits, a subset of which are subjected to experimental testing for the discovery of functionally relevant novel compounds. **b**, Number of phenotypic conditions with at least one positive hit. Numbers above bars indicate the number of perturbations with positive hits over the total number tested. **c**, Hit counts across 11 productive gene perturbations for different compound generation methods. **d**, Distribution of compound–phenotype matching scores for hits and non-hits in the genetic perturbation test set. “Hits” are generated molecules with Tanimoto similarity >0.6 to ground-truth compounds; “no-hits” are dissimilar molecules. Hits exhibit a shift toward higher scores relative to non-hits. **e**, Schematic illustration of the KEAP1–NRF2 signaling pathway. KEAP1 inhibition stabilizes NRF2, allowing its translocation to the nucleus and activation of antioxidant response element (ARE)–driven genes, including *NQO1* and *GCLM*. GEMGen leverages gene expression signatures associated with NRF2 activation to generate candidate KEAP1 inhibitors. **f**, High-scoring molecules (score ≥ 0.6) were ranked by induction levels of ARE-driven genes, revealing an enriched fraction within the top 1% of NRF2 activation compared with randomly sampled L1000 molecules. The bar plot (left) showing the percentage of molecules in the top 1% NRF2 activation percentile for high-score versus random sampling. The dot plot (right) presenting individual generated molecules positioned by their NRF2 activation ranking percentile. **g**, Chemical structure of flavokawain C, a chalcone derivative identified from GEMGen-generated compounds absent from the L1000 dataset. **h**, qPCR analysis of *NQO1* transcript levels in K562 cells treated with different concentrations of flavokawain C. Data are shown as mean ± s.e.m; EC_50_ = 6.3 µM. **i**, Heatmap of RNA-sequencing results showing relative expression levels of NRF2 target genes following treatment with flavokawain C, bardoxolone, or DMSO control. Colors indicate normalized transcript levels. **j**, Western blot analysis of NRF2 protein level in K562 cells following treatment with indicated concentrations of flavokawain C. β-actin was used as loading control.

To this end, we leveraged a CRISPR interference Perturb-seq dataset from Replogle et al.(37), which provides single-cell transcriptomic profiles for genome-scale genetic perturbations. From this dataset, we manually selected 21 genes whose knockdown elicited robust and distinctive transcriptional responses and for which sufficient cell coverage was available. In parallel, we assembled a ground-truth compound set by curating known inhibitors of the selected targets from the ChEMBL database (**Fig. 4a; Methods**).

For 11 of the 21 genetic perturbations, GEMGen generated hit compounds with Tanimoto similarity greater than 0.6 to the corresponding ground-truth inhibitors (**Fig. 4b**), with a variable number of hits across perturbations (**Fig. 4c**). By contrast, existing generative baselines and random sampling from the ChemDiv library showed limited ability to recover ground-truth-like compounds, producing only a single sporadic hit across all perturbation inputs (**Fig. 4b–c**).

We next assessed whether the compound–phenotype scorer could prioritize GEMGen-generated molecules by ranking candidates according to their likelihood of reproducing the target transcriptional signatures. Hit compounds received significantly higher scores than non-hit compounds (**Fig. 4d**), indicating that the scorer is capable of enriching for phenotypically relevant molecules. However, the separation between hit and non-hit score distributions was less pronounced than that observed for the L1000 and Tahoe benchmark datasets. Consistent with this observation, increasing the score threshold did not consistently improve hit rates for several genetic perturbations (**Fig. S6**), suggesting that high-scoring molecules do not always closely resemble known inhibitors.

This discrepancy likely reflects the intrinsic complexity of small-molecule perturbations. Unlike single-gene knockdown, small-molecule inhibitors often induce multifaceted transcriptomic responses due to polypharmacology and off-target effects. As a consequence, high-scoring compounds in this setting do not necessarily resemble known reference inhibitors, raising the possibility that, for specific genetic perturbations, phenotypic matching may arise from chemically distinct compounds operating through alternative mechanisms.

### GEMGen identifies new KEAP1 inhibitors

To explore this possibility in a concrete setting, we focused on *KEAP1*, a genetic perturbation for which score-based prioritization did not consistently recover known inhibitors. KEAP1 is an E3 ubiquitin ligase that targets *NFE2L2* (also known as NRF2) for proteasomal degradation and serves as a central regulator of the cellular oxidative stress response (**Fig. 4e**). Pharmacological activation of NRF2 through KEAP1 inhibition is a validated therapeutic strategy, including for the treatment of Friedreich’s ataxia (38).

Among the compounds generated for the KEAP1 perturbation, 35.2% were present in the L1000 dataset (**Fig. S7**). Leveraging their known transcriptomic profiles from L1000, we evaluated these molecules by assessing NRF2 downstream gene induction and ranking based on their magnitude of induction. As expected, high-scoring molecules (>0.6) ranked above random L1000 compounds (**Fig. 4f; Methods**), demonstrating GEMGen’s ability to recover and prioritize phenotype-recapitulating molecules.

We next examined compounds absent from the L1000 library. Notably, 64.8% of the compounds generated for the KEAP1 CRISPR perturbation signature were not represented in L1000, suggesting that GEMGen can propose molecules beyond existing compound libraries. To determine whether these molecules exhibit *bona fide* KEAP1 inhibitory activity, we prioritized commercially available compounds for experimental validation.

Among the 2,010 GEMGen-generated compounds not present in L1000, 87 were associated with registered CAS numbers. After further filtering based on scorer ranking (score > 0.6) and immediate availability, 16 compounds were selected for experimental testing (**Fig. S8a**). Using the expression of *NQO1*, a canonical NRF2 target gene, as a functional readout, we identified a chalcone derivative, flavokawain C, that dose-dependently induced NQO1 expression in both K562 cells (EC_50_=6.3 µM; **Fig. 4g–h**) and HEK293T cells (EC_50_ = 1.1 µM; **Fig. S8b**).

RNA sequencing followed by gene set enrichment analysis confirmed that flavokawain C upregulated a broad set of NRF2 target genes to a degree comparable to the known NRF2 activator bardoxolone (**Fig. 4i and Fig. S8d**). Gene ontology analysis further supported activation of an antioxidative transcriptional program (**Fig. 4i and Fig. S8c**). Additionally, flavokawain C treatment increased NRF2 protein abundance, suggesting stabilization of NRF2 through reduced KEAP1-mediated ubiquitination and degradation (**Fig. 4j**).

Although chalcone scaffolds have previously been associated with NRF2 activation, likely owing to electrophilic reactivity toward KEAP1 cysteine residues via a Michael acceptor motif, flavokawain C was neither included in the L1000 dataset nor previously reported to exhibit this activity. Therefore, these results demonstrate that GEMGen can phenocopy genetic perturbations by generating chemically novel small molecules with desired phenotypic effects (**Fig. 4a**), highlighting GEMGen’s ability to infer actionable chemical space directly from gene-level perturbations and translate genetic knockdown signatures into functionally relevant small-molecule inhibitors.

### GEMGen generates compounds reverting fibrosis phenotype

To explore the applicability of GEMGen to real-world disease settings, we extended the framework to the transcriptional program underlying fibrosis. Fibrosis comprises a broad spectrum of degenerative disorders characterized by excessive extracellular matrix deposition and tissue scarring across multiple organs, driven by persistent activation of fibroblasts into myofibroblasts. Identifying small-molecule agents capable of reversing or attenuating this cellular program represented a major therapeutic strategy for fibrotic disease. This phenotypic transition is marked by a well-defined transcriptomic signature, including upregulation of *ACTA2*, *COL1A1*, *FN1*, and *TGFB1*. Accordingly, we used this fibrosis-associated transcriptional signature as input to GEMGen to generate candidate compounds predicted to suppress fibroblast activation (**Fig. 5a**).

**Figure 5.**
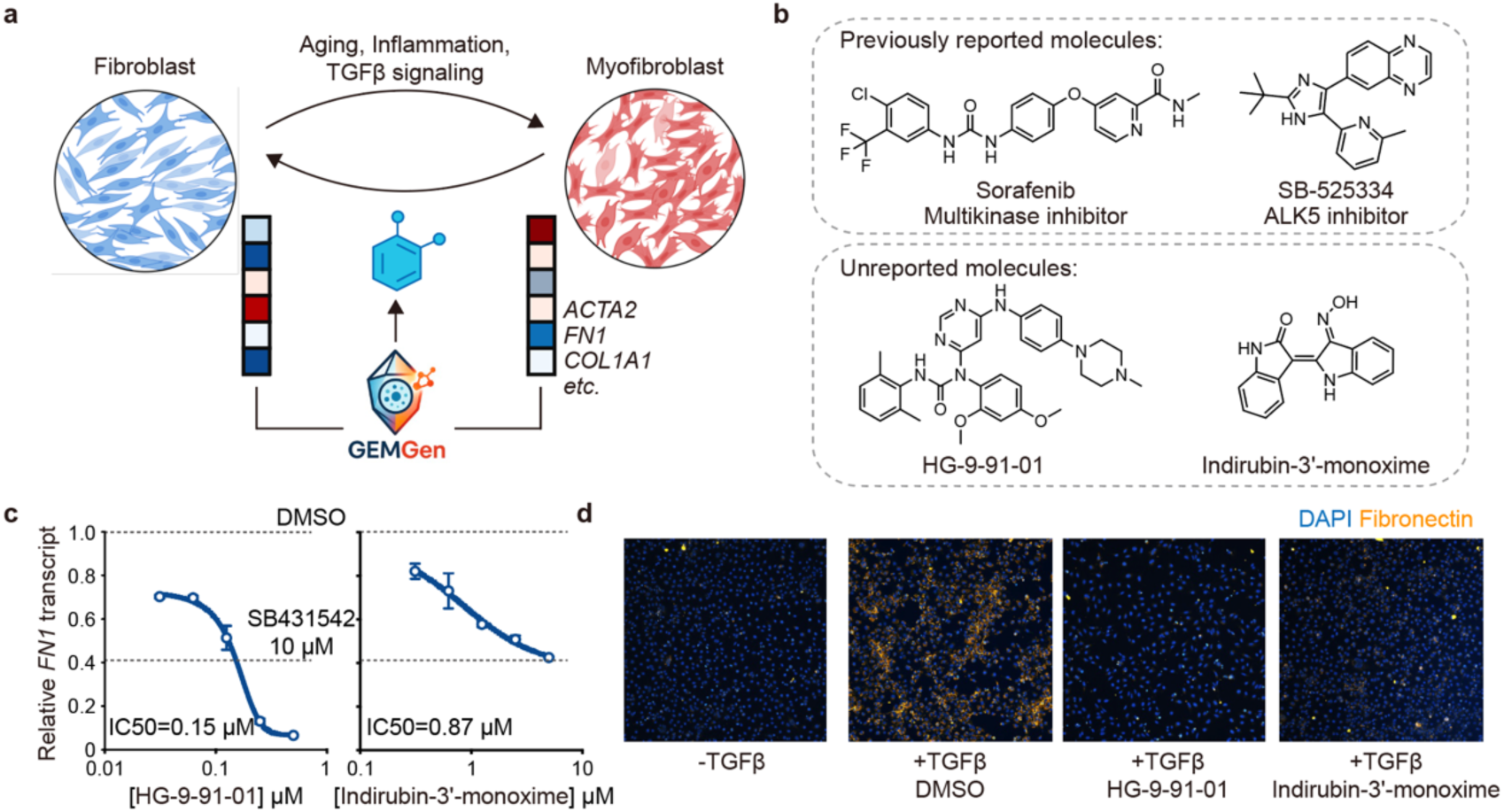
GEMGen generates compounds that reverse the fibrotic fibroblast activation phenotype. **a**, Schematic illustration of fibroblast–myofibroblast state transitions and the application of GEMGen to the fibrosis transcriptional program. Fibroblast activation is associated with upregulation of fibrosis markers including *ACTA2*, *FN1*, and *COL1A1*. The fibrosis-associated transcriptional signature is used as input to GEMGen to generate candidate small molecules predicted to attenuate fibroblast activation. **b**, Chemical structures of representative GEMGen-generated compounds. Upper rows: GEMGen-recovered previously reported molecules. Lower rows: previously unreported molecules selected for experimental validation (HG-9-91-01 and indirubin-3′-monoxime) **c**, qPCR analysis of FN1 transcript levels in TGFβ-stimulated LX-2 cells treated with HG-9-91-01 (left) and indirubin-3′-monoxime (right), normalized to DMSO control levels; SB431542 (10 µM) is included as a positive control. IC50 values are indicated. **d**, Immunofluorescence analysis of fibronectin deposition in fibroblasts treated as indicated. Nuclei are stained with DAPI (blue), and fibronectin is shown in yellow.

Aside from recovering known fibrosis inhibitors such as SB-525334 and sorafenib (**Fig. 5b**), GEMGen prioritized additional candidate molecules predicted to attenuate fibroblast activation. Filtered by annotated bioactivity and commercial availability, we selected 19 molecules for experimental validation. Among these molecules are HG-9-91-01, a salt-inducible kinase inhibitor, and indirubin-3′-monoxime, a dual GSK3β and CDK1/5 inhibitor (**Fig. 5b**). Both compounds suppressed expression of the fibrosis marker *FN1* in TGFβ-stimulated fibroblasts in a dose-dependent manner (**Fig. 5c**). Consistent with transcriptional effects, treatment with either compound markedly reduced TGFβ-induced fibronectin deposition, indicating attenuation of fibroblast activation at the phenotypic level (**Fig. 5d**). Together, these results show that GEMGen-prioritized compounds can functionally revert key features of the fibrotic state by suppressing both fibrosis-associated gene expression and extracellular matrix production. Unlike traditional phenotypic screens, which typically achieve hit rates of only 0.01%–1% due to the need to sceen for large scale chemical libraries, GEMGen dramatically improves efficiency—delivering 2 hits from 19 tested compounds (10.5% hit rate) that potently attenuated TGFβ-induced fibroblast activation (**Fig. 5c–d**). This 10-1000-fold enrichment underscores GEMGen’s power to generate high-confidence candidates, accelerating discovery in complement to conventional physical screening.

## Discussion

In this study, we present an LLM-based generative framework for phenotypic discovery of bioactive molecules that operates directly on transcriptomic state transitions, enabling the proposal and prioritization of candidate perturbations based on desired cellular outcomes rather than predefined molecular targets. By integrating representation learning, generative modeling, and phenotype-based evaluation into a unified system, our approach instantiates a scalable, *in silico* analogue of phenotypic drug discovery that shifts emphasis from static screening toward adaptive exploration of cellular response space.

Viewing disease through a whole-cell, systems-level lens offers fundamental advantages over reductionist paradigms, as it frees therapeutic discovery from reliance on predefined targets or pathways and enables the exploration of non-intuitive biological mechanisms. By operating directly in phenotypic space and vast chemical space, our framework captures coordinated transcriptional state changes that reflect integrated cellular responses, rather than isolated molecular perturbations. This perspective allows the identification of candidate interventions that would be difficult, if not impossible, to uncover when starting from a single target hypothesis.

The framework exhibits substantially expanded generative capacity compared with prior phenotype-based computational models, such as DrugReflector (16), which primarily focus on embedding, retrieval, or ranking of existing compounds. By explicitly learning a forward generative process over transcriptional perturbations, our model proposes novel candidates, which extend beyond interpolation within known chemical space, as starting points for experimental validation and molecular optimization, potentially yielding therapeutic candidates.

More broadly, this work demonstrates how cell-state–level generative modeling can serve as a practical instantiation of systems-level therapeutic discovery. By operating on coordinated gene expression programs rather than isolated molecular interactions, the approach aligns naturally with emerging views of disease as a perturbation of cellular state landscapes (39).

In addition, a central strength of this work lies in the closed-loop coupling of generation and evaluation. This generative–evaluative loop enables efficient traversal of a high-dimensional biological response landscape and facilitates the discovery of non-intuitive interventions that may be missed by target-centric or library-bound approaches.

Several limitations of the present study should be acknowledged. First, the framework relies on the availability and quality of transcriptomic perturbation data, which may incompletely capture context-specific cellular responses or long-term phenotypic consequences. Second, while transcriptomic similarity provides a powerful proxy for phenotypic alignment, it does not directly encode downstream functional, physiological, or safety-related outcomes, which will require integration of additional modalities in future extensions. Finally, although the generative model expands accessible candidate space, experimental validation remains essential to confirm biological efficacy and translational relevance.

Addressing these limitations will require the incorporation of multi-omic and functional readouts, improved modeling of cellular context and dynamics, and tighter integration with experimental feedback. Nevertheless, the framework introduced here provides a foundation for scalable, adaptive phenotypic discovery and illustrates the potential of generative models to reshape how therapeutic hypotheses are formulated and explored.

## Methods

### GEMGen Training

GEMGen is fine-tuned from NatureLM for phenotype-based compound generation and scoring. The training was implemented through three complementary instruction–response tasks.

#### NatureLM foundation model

NatureLM (28) is a unified scientific language model trained on heterogeneous data modalities, including biological sequences (e.g., protein FASTA and molecular SMILES), natural language text, and other cross-modal scientific data. It adopts a decoder-only Transformer architecture, analogous to GPT, and is trained autoregressively by representing all modalities as token sequences within a shared vocabulary.

NatureLM is released in multiple model sizes. Given the limited scale of cell–compound perturbation data available for our downstream tasks, and to reduce the risk of overfitting to compound-specific patterns, we employed the 1B-parameter variant reproduced by Zhang et al. (40). This model consists of 16 Transformer layers with a hidden size of 2,048, a feed-forward network (FFN) dimension of 5,504, and 32 attention heads, including 8 key–value (KV) heads. The vocabulary comprises 130,239 tokens spanning diverse scientific entities and English text, including 1,401 tokens corresponding to unique SMILES symbols. As a general-purpose scientific foundation model, NatureLM provides a strong and flexible initialization for text-guided molecular generation and compound–phenotype modeling tasks.

#### Bioentity representation task

This task was implemented through instruction fine-tuning of NatureLM on a diverse mixture of biological data. Descriptive annotations were curated for a wide range of human and mouse cell types and protein-coding genes (detailed data preparation procedures are provided in subsequent sections). Each entry was reformatted into an instruction–response structure with the assistance of GPT-4o. This process yielded 255 unique instruction–response pairs for cell type descriptions and 75,203 for gene descriptions. During training, the two data types were combined and randomly shuffled while maintaining their original distribution ratios. The model was trained using a cross-entropy loss function, consistent with the optimization setup used in NatureLM.

#### Phenotype-guided compound generation task

The L1000 chemical perturbation transcriptomic dataset (12), containing paired compound and perturbed-transcriptome information, was used to train the model for phenotype-guided compound generation task. Transcriptomic signatures were transforms into text-based ranked lists of differentially expressed genes as input. Training was performed in two phases to balance data coverage and model performance. Phase I was conducted on compound perturbations curated from the original L1000 dataset. The model was trained for two epochs across 16 GPUs using a cross-entropy loss function, a learning rate of 1e-5, and a batch size of 256. Phase II involved continued instruction fine-tuning with the L1000CDS² dataset (41), which provides higher-quality measurements and broader gene coverage. This stage was designed to improve generalization across diverse transcriptomic profiling platforms and protocols. Training in Phase II was carried out for 2,475 iterations on 8 GPUs with a cross-entropy loss function, a reduced learning rate of 8e-6 and a batch size of 64. Details of data preprocessing and train/test splitting are provided in subsequent sections.

#### Compound–phenotype matching task

A separate model was fine-tuned to specialize in scoring the correspondence between chemical perturbations and transcriptomic changes, represented as text-based ranked lists of differentially expressed genes. Similar to the compound generation task, training was conducted in two sequential phases using the same data corpus. To adapt the framework for scoring, several task-specific modifications were introduced.

The model was trained to produce a binary response (“yes” or “no”) for each compound–phenotype pair, indicating whether the compound was predicted to reproduce the given transcriptomic change. The training objective combined two complementary losses: a cross-entropy loss for autoregressive language modeling and a binary cross-entropy loss applied to a classification head. Specifically, the final hidden state corresponding to the response token (“yes” or “no”) was passed through a two-layer multilayer perceptron (MLP; hidden dimension = 2,048) for explicit binary classification. During inference, we use the output from the classification head as the final binary prediction.

For scorer training, Phase I was conducted for two epochs on 8 GPUs with a learning rate of 2e-5 and a batch size of 512. Phase II continued for 3,008 steps on 8 GPUs using a reduced learning rate of 8e-6 and a batch size of 128.

### Datasets

#### L1000 dataset

We retrieved the Level 3 transcriptomic profiles and associated metadata from the L1000 dataset of the CMap LINCS (Library of Integrated Network-Based Cellular Signatures) resource (12) (https://clue.io/data/CMap2020#LINCS2020, December 2020 release). In our study, we focused exclusively on compound perturbation profiles. The Level 3 data contain quantile-normalized expression values for 978 landmark genes (experimentally measured) and an additional 11,350 genes computationally inferred. For our task, we used only the measured expression values of the 978 landmark genes.

To compute differential gene expression (DGE) signatures, we applied the Characteristic Direction (CD) method (42). CD is a multivariate geometrical approach that quantifies expression change levels and computes statistical significance for each gene between two groups of transcriptomic profiles. Specifically, we grouped samples based on perturbation metadata—including cell type, compound, time, dose, and plate ID. Samples sharing identical perturbation conditions on the same plate were considered replicates. We computed DGE signatures for each such group against the matched vehicle-treated controls on the same plate to minimize confounding from plate-specific technical variability.

For each cell type–compound pair (hereafter referred to as a combination), multiple transcriptomic experiments may be available, conducted under different time points, doses, or plates. We refer to all such experiments within a given combination as replicates, regardless of the specific experimental settings. To ensure data quality and prevent overrepresentation of certain combinations, we retained two DGE signatures per combination, selected based on internal consistency measured by cosine similarity:

- For combinations with 2 to 20 replicates (54.6% of all combinations), we computed pairwise cosine similarities among all DGE signatures and selected the two with the highest average similarity to the rest.
- For combinations with more than 20 replicates (1.2%), we first applied Leiden clustering to group similar DGE signatures, then calculated intra-cluster cosine similarities. The two most consistent DGE signatures were selected from the cluster with the highest mean internal similarity.
- For combinations with only one replicate (44.2%), we generated five synthetic control sets by repeatedly sampling control cells (matched in size to the perturbed group) from the same plate. We computed five CD-based DGE signatures and selected the two with the highest pairwise similarity.

This procedure yielded a curated dataset of 183,440 unique cell type–compound combinations, each represented by two high-consistency DGE signatures, spanning 27,770 compounds and 213 cell types.

#### L1000CDS^2^ dataset

L1000CDS^2^ is a publicly available database of precomputed CD-derived DGE signatures derived from the L1000 dataset (41). We downloaded 119,155 DGE signatures from the official L1000CDS^2^ database (https://maayanlab.cloud/public/L1000CDS_download/), and filtered entries with compound perturbations and adjusted *p*-values less than 0.1, resulting in 32,809 high-confidence records. These records span 63 human cell lines and 3,872 small molecules.

Compared to the CD-derived DGE signatures we computed from the L1000 dataset, the L1000CDS^2^ signatures include both landmark and inferred genes, thereby offering broader transcriptomic coverage. This expanded gene space is particularly valuable for model training in scenarios where the downstream applications (e.g., single-cell or other non-plate-based sequencing data) are not restricted to landmark genes alone.

#### Train/test split

To evaluate the model’s ability to generalize to previously unseen chemical space, we constructed a novel compound test set based on the L1000 and L1000CDS^2^ datasets. The key criterion was that no compound in this test set exhibits a Tanimoto similarity greater than 0.8 to any compound used during training.

To achieve this, we first pooled all small molecules from the filtered L1000 set and the L1000CDS² dataset into a unified compound collection and computed the pairwise Tanimoto similarity matrix across all molecules. For each compound in L1000CDS², we identified its maximum similarity to any other compound in the combined set. Compounds whose maximum similarity was below 0.8 and that originated from L1000CDS² were retained as candidates for the novel compound test set. The resulting novel compound test set comprises 404 samples.

#### Tahoe-100M dataset

The recently released Tahoe-100M dataset (29) captures the transcriptional responses of 50 cancer cell lines to 335 compound perturbations using the Mosaic platform. We downloaded the full dataset and associated metadata from the official Tahoe-100M repository hosted on Hugging Face [https://huggingface.co/datasets/tahoebio/Tahoe-100M].

For data preprocessing, we applied the following strategy: First, cells were filtered using quality-control criteria, including a mitochondrial gene fraction below 20%, more than 50 expressed genes, and restriction to the G1 phase of the cell cycle.

Second, only cells treated with a 5 μM drug dose were retained to ensure consistency across perturbation conditions. Third, for each cell type–compound combination, we randomly sampled 100 cells before and 100 cells after perturbation. Fourth, we computed the E-distance between perturbed and control cells (43), which quantifies the effect size of perturbation in high-dimensional expression space. We retained combinations with E-distance > 500, resulting in 640 high-confidence cell type–compound combinations with strong perturbation effects. Finally, we calculated CD-derived DGE signatures for each selected combination for model evaluation.

#### K562 CRISPR perturbation dataset

The CRISPR perturbation dataset for the K562 cell line used in this study was derived from the experiments conducted by Replogle et al (37). We downloaded the scRNA-seq dataset of the K562 cell line before and after different gene edits from the website https://gwps.wi.mit.edu/. Following the analysis described in the original paper, we selected 21 genetic perturbations that induce significant transcriptomic changes and identified the corresponding sets of upregulated and downregulated genes. Under the assumption that chemical inhibition of a protein phenocopies the effects of genetic knockdown or knockout, we manually curated a list of small molecule inhibitors of each target from the ChEMBL database, restricting to those characterized by quantitative, protein-based assays. To preclude low-activity inhibitors, only high-potency inhibitors with IC50 values below 500 nM were retained as ground-truth molecules for analysis.

#### Cell type descriptive annotations

We curated descriptive annotations for all cell types used in this study by combining two sources: descriptions retrieved from the ATCC cell catalog, and independently generated summaries by GPT-4o, which were manually reviewed to ensure biological plausibility. In total, we compiled 255 cell type descriptions, including 42 primary cells and 213 immortalized cell lines.

#### Gene descriptive annotations

We retrieved human and mouse protein-coding gene list from the NCBI database (https://ftp.ncbi.nlm.nih.gov/gene/DATA/; version dated April 2025). For each gene, we extracted descriptive summaries from the RefSeq database using the official API. In total, we collected annotations for 20,786 human genes and 18,756 mouse genes.

### Instruction-response pair construction

#### phenotype-compound pair

To support both training and inference for the compound generator and the scoring module, we constructed instruction–response pairs derived from phenotype-compound pairs.

From each DGE signature, we derive upregulated and downregulated gene list based on the CD vector, which encodes the magnitude and direction of gene differential expressions. Specifically, we selected the top 50/40/30 genes with the largest absolute CD values and then assigned each gene to the upregulated or downregulated category based on the sign of its value (positive or negative). This yielded two directional gene sets per DGE signature.

Given that many small molecules appear across multiple cell type–compound combinations, we introduced molecular representation diversity by generating alternative SMILES strings for each compound using the SMILES augmentation method implemented in TamGen (26) [https://github.com/microsoft/TamGen]. This augmentation enriches the chemical space explored during generative modeling and helps prevent overfitting to SMILES forms in the training dataset.

For both the generator and the scorer, the input and output take the form of natural language instructions and responses, following a standardized prompt-response interface as shown in Figure S1. This unified format allows both models to interface with natural language prompts in instruction-based generation and evaluation workflows.

#### Cell type and genes

To construct instruction–response pairs for cell types, we first manually designed a base instruction template (for example, *“Describe the cell type [cell type]”*). To increase linguistic diversity and reduce structural uniformity across instructions, we then used GPT-4o to generate paraphrased variants of this template (for example, we prompted GPT-4o with: *“Here is a prompt for generating cell type descriptions: ‘Describe the cell type [cell type].’ Please generate 20 similar sentence templates based on this example.”*). For each cell type, we randomly selected one instruction from the augmented template pool. The corresponding response was the descriptive annotation of the cell type, as curated from ATCC and GPT-4o (see the previous section).

We followed a similar procedure for genes, except that we generated 50 instruction templates in total. Additionally, because gene names often have multiple synonyms, we incorporated those synonyms when constructing instruction–response pairs. This resulted in a total of 75,203 distinct instruction–response pairs for gene entities.

The full list of instruction templates is provided in **Table 1**.

**Table 1.**
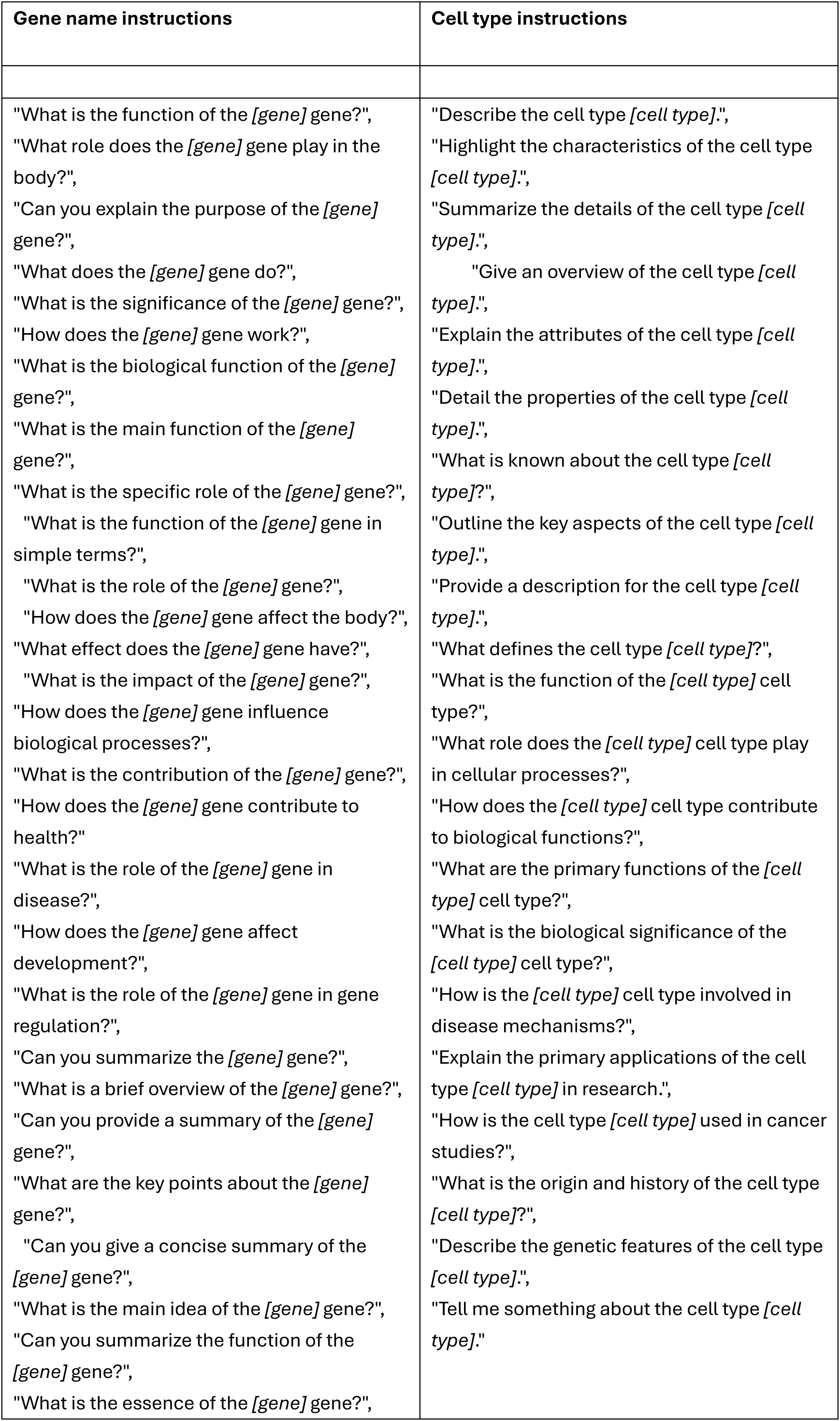
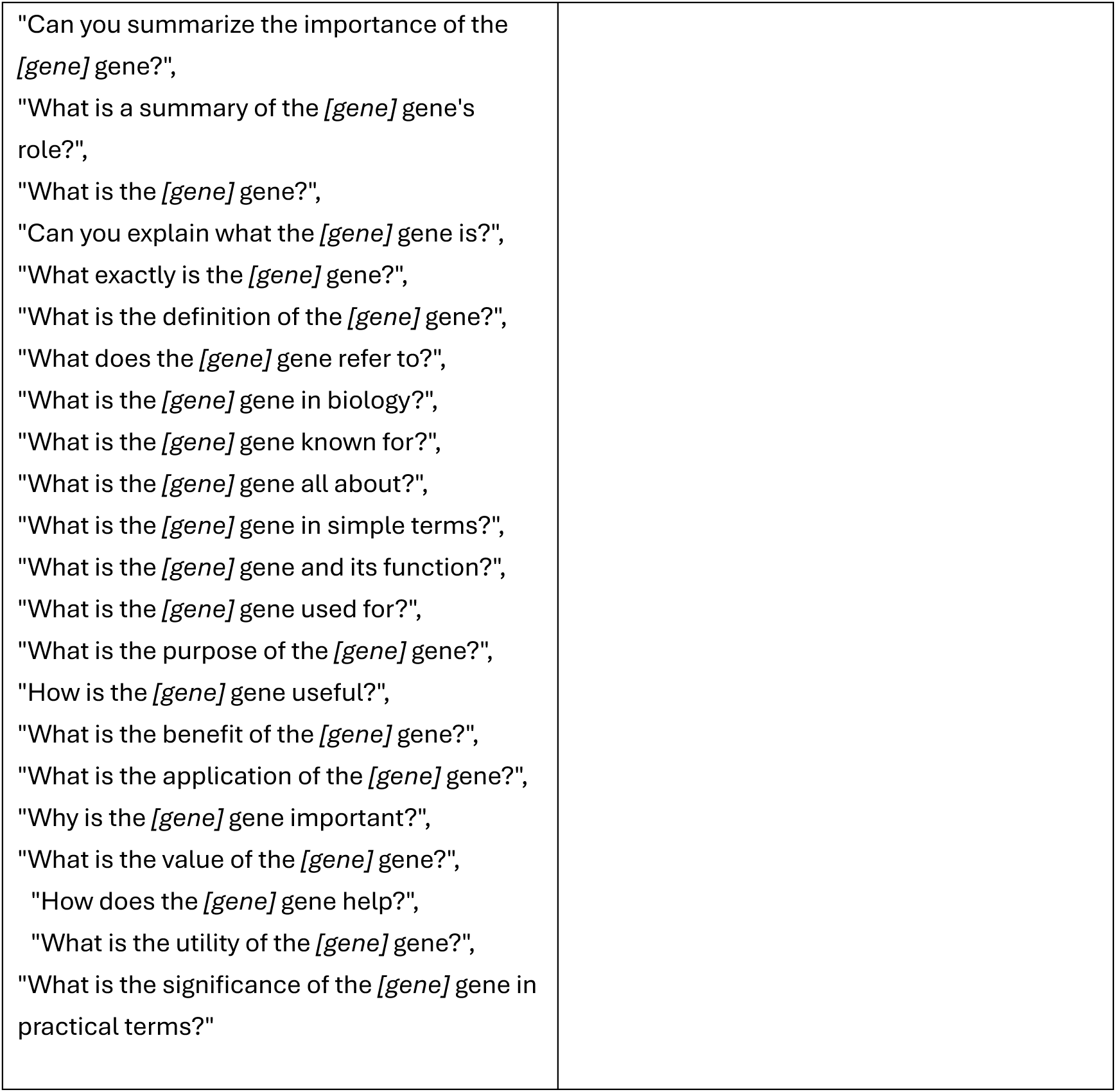
GPT-4o augmented instruction templates.

### Cross-Dataset Consistency Evaluation

To evaluate the ability of text-based representations to capture cross-dataset consistency in drug perturbation responses, we compared conventional numerical differential expression analysis with GEMGen’s language-model–derived representation learning. We first identified overlapping cell line–drug perturbation pairs shared between the L1000 and Tahoe100M datasets, yielding 108 matched pairs. For each pair, expression profiles of 933 genes common to both datasets were extracted. Within each dataset, control samples from the same cell line were randomly assigned to each perturbed condition. For the numerical baseline, differential expression analysis was performed using the Wilcoxon rank-sum test implemented in Scanpy to compute log₂ fold-change values. Cosine similarity was then calculated between the log₂ fold-change vectors derived from the two datasets for each matched perturbation pair. In parallel, we applied the CD method to identify differentially expressed genes within each dataset. The resulting gene sets were combined with cell line information to construct textual prompts, which were subsequently processed by GEMGen to obtain sentence embeddings. Cosine similarity was then computed between sentence representations derived from the two datasets for each matched pair. To account for methodological differences between the numerical and text-based approaches, negative control pairs were generated by randomly shuffling the matched perturbation combinations. Finally, kernel density estimates of cosine similarity distributions for positive and negative pairs were compared under both computational frameworks.

### Negative Sample Construction for scorer training

By design, the compound–phenotype scoring model functions as a binary classifier, predicting whether a given small molecule is consistent with a specified DGE signature in a particular cell type. However, the observed cell–compound perturbation data only provide positive samples—pairs in which the compound is known to induce the given transcriptional response. To enable effective binary training, we constructed negative samples using a similarity-based replacement strategy.

For each positive sample, we generated a matched negative example by replacing the compound with a structurally dissimilar molecule. Specifically, we randomly selected the replacement compound from the 100 compounds in the training set with the lowest Tanimoto similarity (based on Morgan fingerprints) to the original compound. All other components, including the cell type and DGE signature, were kept unchanged, and the sample was relabeled as negative.

This strategy ensures that the resulting negative examples are chemically plausible yet biologically implausible, enabling the model to learn to distinguish true perturbation–response relationships from incompatible pairs. The final dataset was balanced, containing one negative sample for each positive example.

### Randomly paired compound–phenotype samples for scorer evaluation

To assess GEMGen’s performance, we constructed randomly paired compound–phenotype control samples and compared their scores with those of non-hit molecules, defined as generated compounds with Tanimoto similarity below 0.6 to ground-truth compounds. For this purpose, non-hit compounds associated with each test phenotype were replaced through the following procedure. First, we randomly selected candidate molecules from the pool of valid compounds generated for other test samples. To ensure structural dissimilarity and avoid trivial matches, candidate replacements were required to satisfy a structural constraint: the maximum Tanimoto similarity between any replacement molecule and the set of molecules originally generated for the target phenotype had to be below 0.4. The selected replacement compound was then paired with the target phenotype to form a randomly matched control sample.

### Baselines

#### Gex2SGen (18)

Since the original implementation of Gex2SGen was not publicly available, we reconstructed the model based on the architectural and training specifications described in the publication. We implemented two variational autoencoders: SMILES-VAE employs an encoder-decoder framework with two stacked GRU layers for processing chemical molecular representations, while p-VAE utilizes a multilayer perceptron architecture to learn latent representations of transcriptomic differential expression profiles, with both latent spaces dimensioned to 256. During pre-training, SMILES-VAE was trained on 1.86 million molecules from the ChEMBL and L1000 databases, and p-VAE was trained on 388,466 differential expression profiles from the L1000 project. To accommodate transcriptomic data from different test sets, we identified the gene intersection between the L1000 and Tahoe100M datasets, comprising a total of 933 genes. For all molecules involved in pre-training, we filtered them using the maximum sequence length of 208 from the test set as the threshold. In the joint training phase, we utilized 388,466 paired transcriptomic expression profiles and molecular SMILES data, connecting p-VAE’s encoder and SMILES-VAE’s decoder through a 256-dimensional adapter layer to enable cross-modal generation from gene expression data to molecular structures. The model’s loss function combines reconstruction loss with KL divergence, employing a progressive weighting strategy for KL terms to ensure latent space continuity and generation diversity. During testing, differential expression vectors from the test set are used as input to generate 1,000 molecular structures through repeated sampling.

#### GexMolGen (17)

Following the instructions provided in the code repository of GexMolGen, we downloaded and installed the corresponding code package and model weights. To accommodate the omics input requirements of GexMolGen, we performed corresponding data processing for each test set. For the L1000CDS^2^ test set, we identified the corresponding pre- and post-perturbation transcriptomic data in the L1000 database based on cell line and perturbation information. Where multiple replicate experiments existed, we averaged their values, utilizing only the 978 truly measured landmark genes. For the Tahoe100M and K562 test sets, we identified cells before and after each perturbation based on the perturbation information. Through sampling, we ensured equal numbers of control and perturbed groups, then computed their respective averages. In this case, we utilized the top 2000 highly variable genes. During testing, we processed one sample at a time, setting the sampling parameter *beam_width* to 1000 and the generation parameter greedy to False. This configuration ensured the generation of 1000 small molecules per sample while preventing excessive repetition in the generated molecular structures.

#### ChemDiv

We downloaded the entire set of 1.53 million small molecules from ChemDiv’s o]icial website (https://www.chemdiv.com/) and employed a random sampling with replacement method to generate 1,000 small molecules per test sample as the output of the virtual screening.

#### GPT-5

GPT-5 is a widely used general-purpose large language model with strong cross-domain reasoning and generation capabilities. We adopted its dialogue-optimized variant, *gpt-5-chat-latest*, as a baseline due to its reliable instruction-following behavior and stable structured outputs. To ensure a fair and controlled comparison, the prompt provided to GPT-5 was kept identical to that used for the fine-tuned GEMGen model. To better elicit the capabilities of gpt-5-chat-latest under our task setting, we designed the following system instruction:

> *“You are a SMILES generator. Your only task is to generate exactly one valid SMILES string of drug-like small molecules, based on a provided panel of genes that are either up-regulated or down-regulated, listed in order of regulation magnitude. The generated SMILES string must have the potential to induce the specified gene panel changes.*

> *Instructions:*

- *Each molecule must be wrapped individually in <mol> and </mol> tags*.
- *Do not include any explanations or text outside the <mol></mol> tags*.
- *Output exactly one molecule — no more, no less.”*

This strict format specification ensured comparable and structurally constrained baseline outputs. For each input phenotype, the instruction was executed 1,000 times to generate candidate molecules.

### Code availability

GEMGen is available on GitHub (https://github.com/DLS5-Omics/GEMGen).

### Experiments

#### Dose-Response Analysis of Flavokawain C

K562 cells were cultured in RPMI 1640 medium (Corning, 10-040-CV) supplemented with 10% fetal bovine serum (CellMax, SA211.02). HEK293T cells were cultured in high glucose DMEM medium (Corning, 10-013-CV) supplemented with 10% fetal bovine serum (CellMax, SA211.02). Cells were treated with flavokawain C (TargetMol) for 24 hours at final concentrations of 40 μM, 20 μM, 10 μM, 5 μM, 2.5 μM, 1.25 μM, and 0.6125 μM. Cells treated with DMSO were included as the control group. At study end, total RNA was extracted from the cells (TRIzol, 15596018CN, ThermoFisher) and reverse-transcribed into cDNA (Takara, RR047A). Quantitative PCR (qPCR) (Takara, RR820) was performed to measure the mRNA expression level of *NQO1*. The dose-dependent effects of flavokawain C on *NQO1* expression were analyzed in both K562 and 293T cells.

#### RNA-seq

K562 cells were cultured in 10-cm dishes and treated with flavokawain C or the known NRF2 activator bardoxolone at a final concentration of 10 μM or 100 nM, respectively. Cells treated with DMSO served as the control group. After 24 hours of treatment, total RNA was extracted using TRIzol reagent according to the manufacturer’s instructions. The purified RNA samples were submitted to GeneWiz (Genewiz) for RNA sequencing. The resulting RNA-seq data were subjected to downstream bioinformatics analysis to evaluate gene expression changes induced by flavokawain C and bardoxolone.

#### Western Blot

HEK293T cells were seeded into 6-well plates and treated with flavokawain C at final concentrations of 5, 10, and 20 μM for 24 hours. Cells treated with DMSO were used as the control group. After treatment, cells were harvested and total cellular protein was extracted using RIPA lysis buffer (Millipore, 20-188). Protein samples were subjected to SDS–PAGE and transferred to membranes for Western blot analysis. Membranes were blocked with 5% BSA in TBST for 1 hour followed by incubation with primary antibodies (NRF2: proteintech, 80593-1-RR, β-actin: proteintech, 66009-1-Ig) in 5% BSA/TBST. Secondary antibodies (Sigma-Aldrich, A9044-2ML, A0545-1ML) were diluted in TBST. NRF2 protein levels were detected, and β-actin was used as the internal loading control.

#### Dose-Response Analysis of HG-9-91-01 and Indirubin-3’-Monoxime

LX-2 cells were seeded into 12-well plates and cultured in DMEM supplemented with 10% FBS for 12 hours. Cells were then treated with HG-9-91-01 or indirubin-3′-monoxime (TargetMol) at final concentrations of 40 μM, 20 μM, 10 μM, 5 μM, 2.5 μM, 1.25 μM, and 0.6125 μM in DMEM containing 0.1% FBS for 12 hours. 10 µM SB431542 and DMSO was used as control groups. Without changing the medium, TGF-β (PeproTech, AF-100-21C-2) was directly added to the culture medium at a final concentration of 10 ng/mL, and cells were further incubated for 48 hours. At study end, total RNA was extracted from the cells followed by qPCR to measure *FN1* mRNA expression, with *ACTB* used as the internal reference gene.

#### Immunofluorescent staining

LX-2 cells were seeded into 12-well plates and cultured in DMEM supplemented with 10% FBS for 12 hours. Cells were then treated with 10 µM SB431542, 200 nM HG-9-91-01, or 3 µM indirubin-3′-monoxime (TargetMol) in DMEM containing 0.1% FBS for 12 hours followed by stimulation with 10 ng/mL TGF-β for 48 hours. At study end, cells were fixed with a 4% paraformaldehyde solution for 10 min and subsequently permeabilized with 0.5% Triton X-100 in PBS followed by blocking for 1 h with PBS containing 0.1% Triton X-100 and 5% FBS and then staining overnight at 4 °C with primary anti-fibronectin antibody solution (proteintech, 66009-1-Ig, 5% FBS, 0.1% Triton X-100 in PBS). Cells were then stained with a secondary antibody solution containing Hoechst 33342 dye (10 μg/mL) for 1 h at room temperature in the dark. Cells were then washed three times with PBS prior to imaging.

---

**Figure S1.**
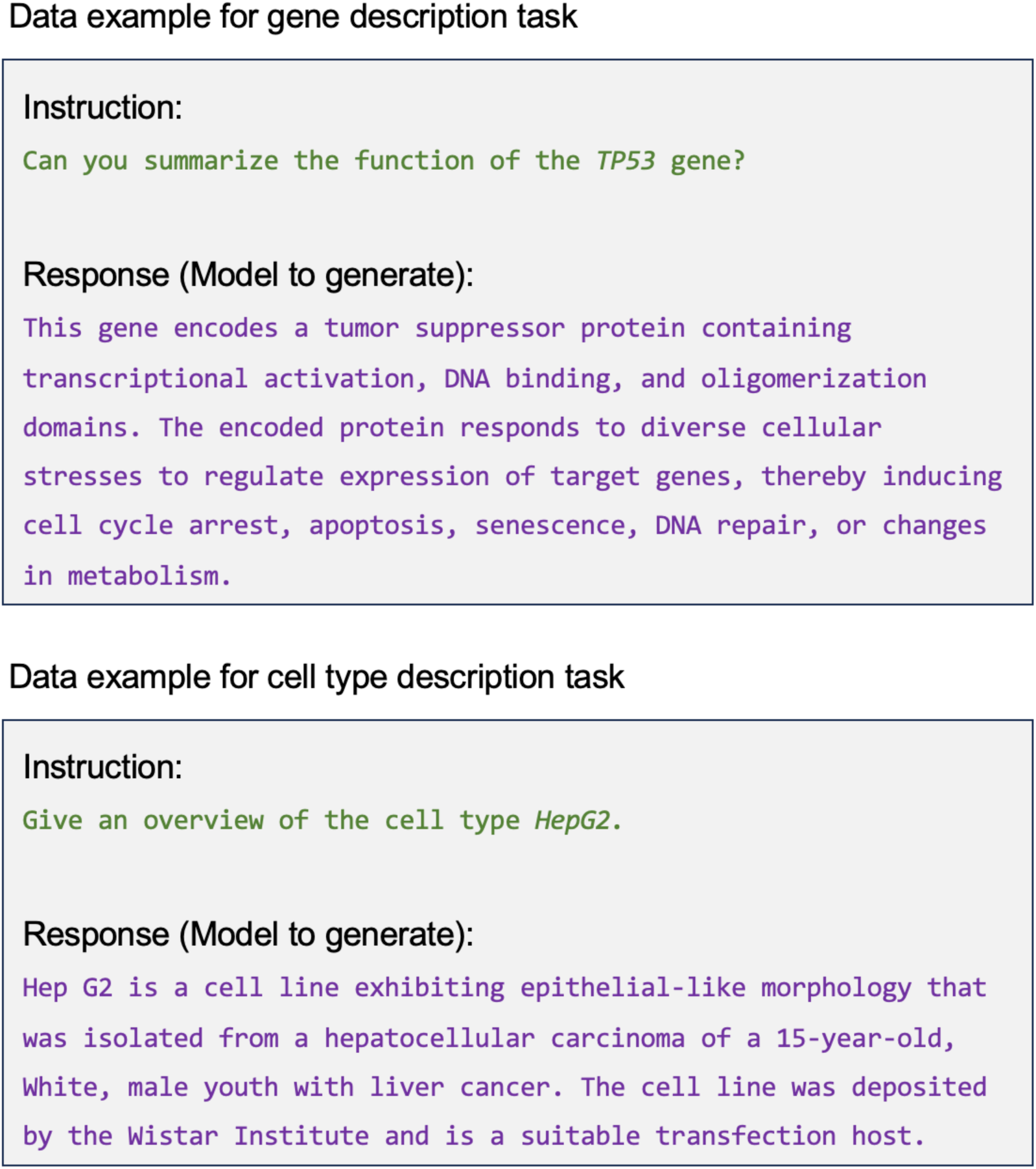
Data examples for bioentity representation generation tasks.

**Figure S2.**
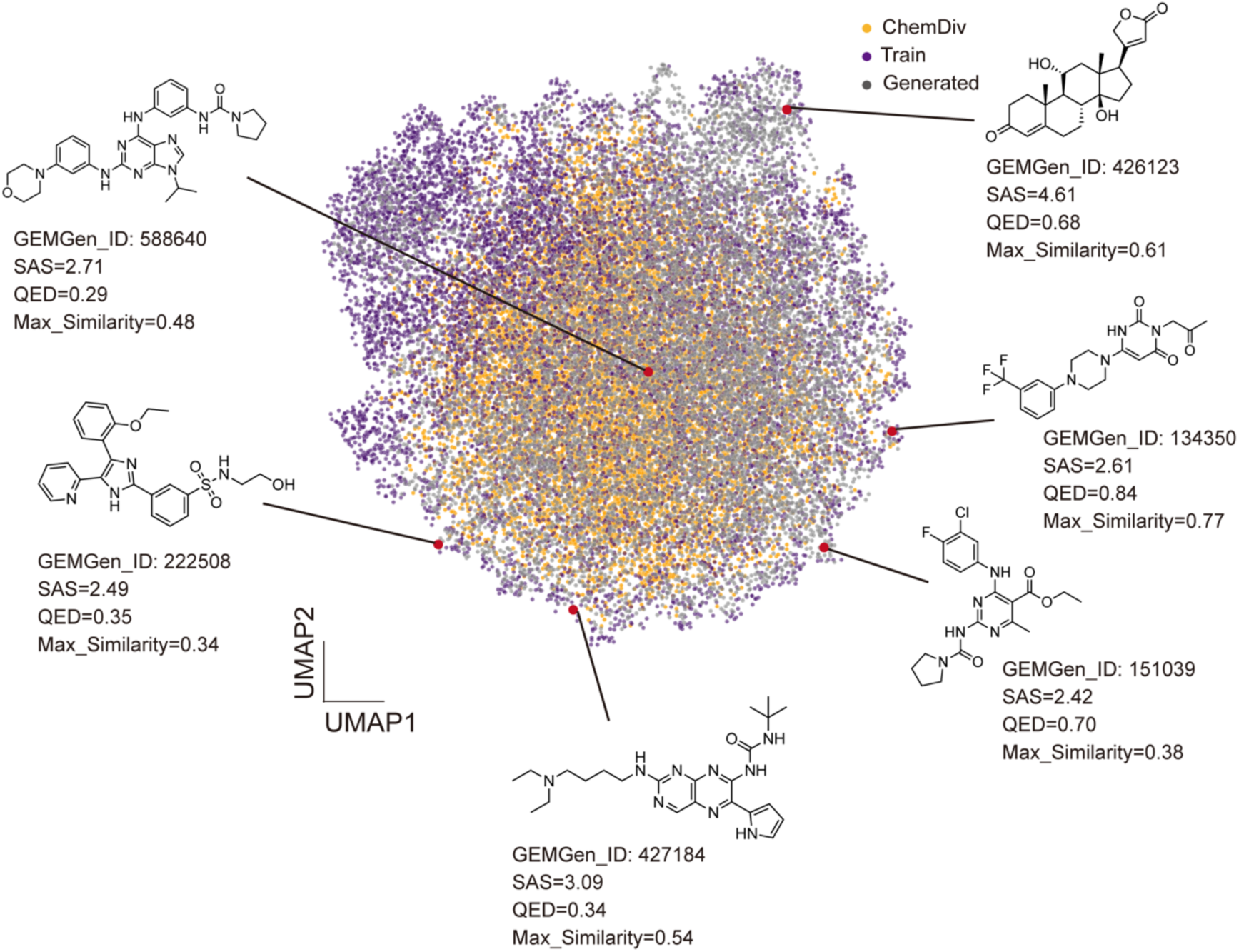
Representative novel molecules generated by GEMGen. UMAP visualization of GEMGen-generated compounds (grey), training compounds (purpue), and ChemDiv compounds (yellow), with representative novel molecules generated by GEMGen. Quantitative analyses of their SAS score, QED, and structural similarity to the training set are provided.

**Figure S3.**
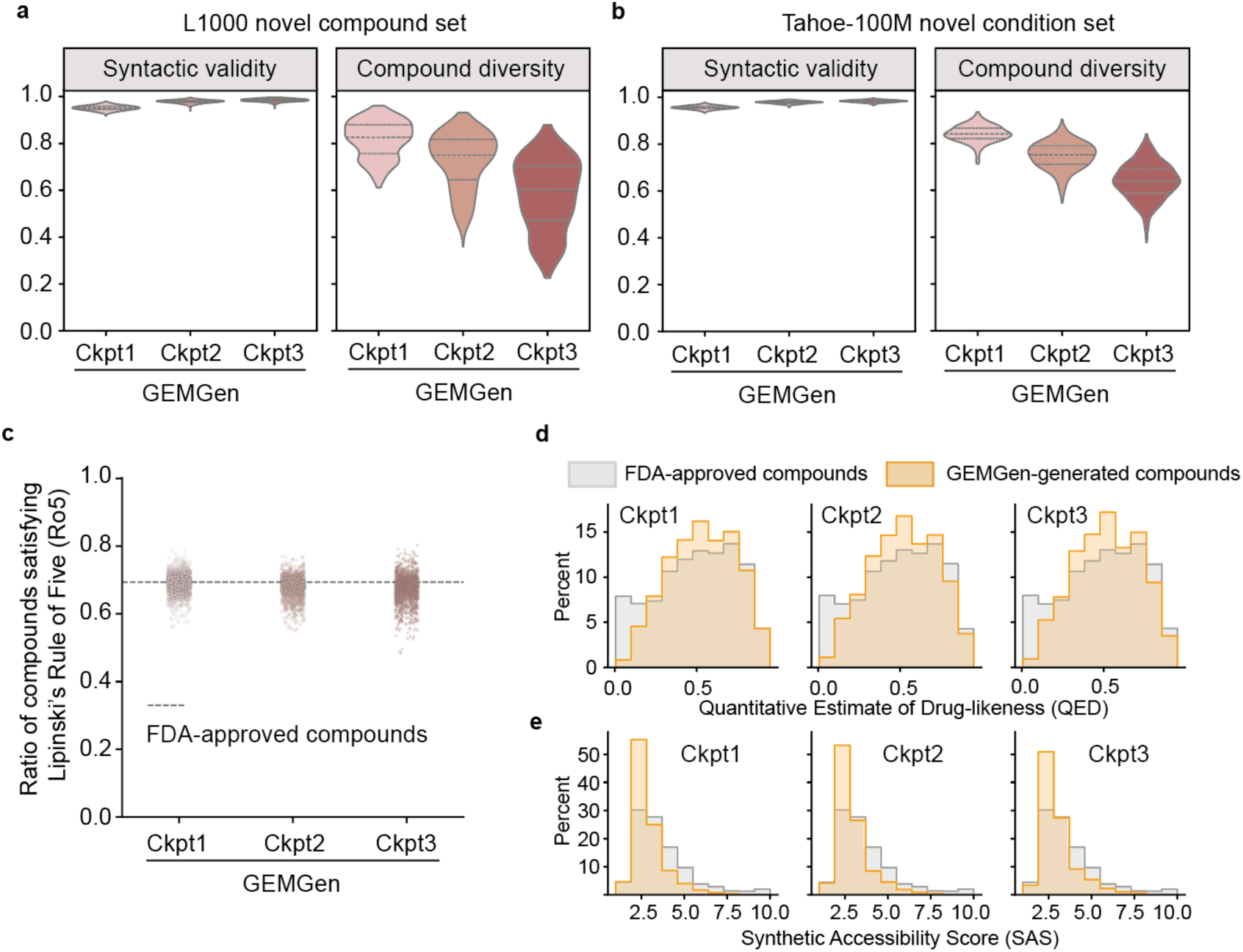
Quality and chemical diversity of compounds generated by GEMGen. **a–b**, Syntactic validity and compound diversity of molecules generated by three GEMGen variants (ckpt1–3), trained for increasing numbers of epochs, on the **(a)** L1000 novel compound test set (*n* = 404) and **(b)** Tahoe-100M novel condition test set (*n* = 640). Syntactic validity measures the fraction of chemically valid SMILES strings among all molecules generated for each input sample. Compound diversity measures the number of unique valid molecules generated per sample. Dashed lines: the 25^th^, 50^th^, and 75^th^ percentiles. **c**, Fraction of generated compounds satisfying Lipinski’s Rule of Five (Ro5) compared to FDA-approved compounds (dashed line). **d**, Distributions of the quantitative estimate of drug-likeness (QED) for generated and FDA-approved compounds. e, Distributions of the synthetic accessibility score (SAS) for generated and FDA-approved compounds. Lower SAS indicates easier chemical synthesis.

**Figure S4.**
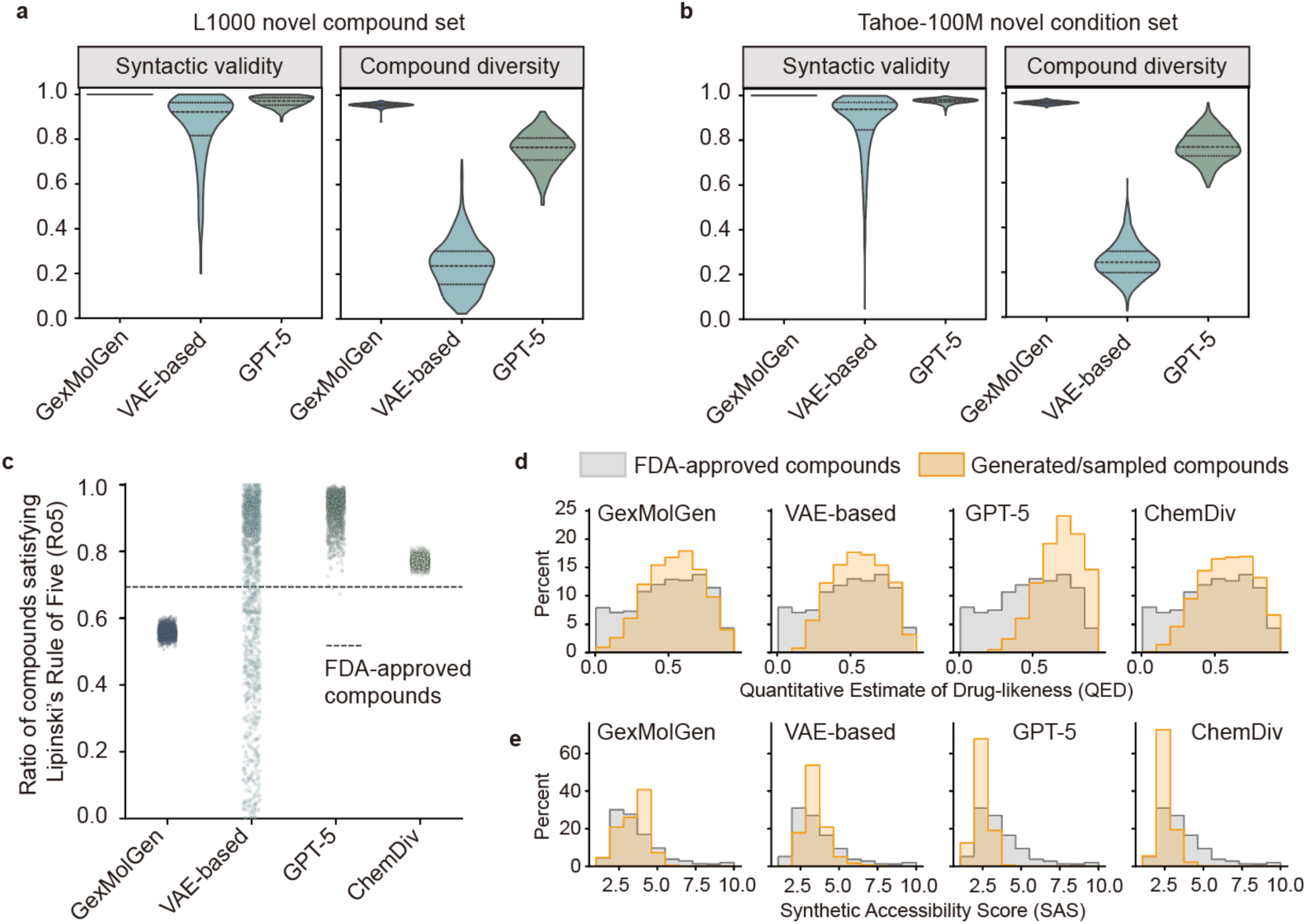
Quality and chemical diversity of compounds generated by alternative approaches. **a–b**, Syntactic validity and compound diversity of molecules generated by GexMolGen, VAE-based method, and GPT-5, on the **(a)** L1000 novel compound test set (*n* = 404) and **(b)** Tahoe-100M novel condition test set (*n* = 640). Syntactic validity measures the fraction of chemically valid SMILES strings among all molecules generated for each input sample. Compound diversity measures the number of unique valid molecules generated per sample. Dashed lines: the 25^th^, 50^th^, and 75^th^ percentiles. **c**, Fraction of generated compounds satisfying Lipinski’s Rule of Five (Ro5) compared to FDA-approved compounds (dashed line). **d**, Distributions of the quantitative estimate of drug-likeness (QED) for generated and FDA-approved compounds. e, Distributions of the synthetic accessibility score (SAS) for generated and FDA-approved compounds. Lower SAS indicates easier chemical synthesis.

**Figure S5.**
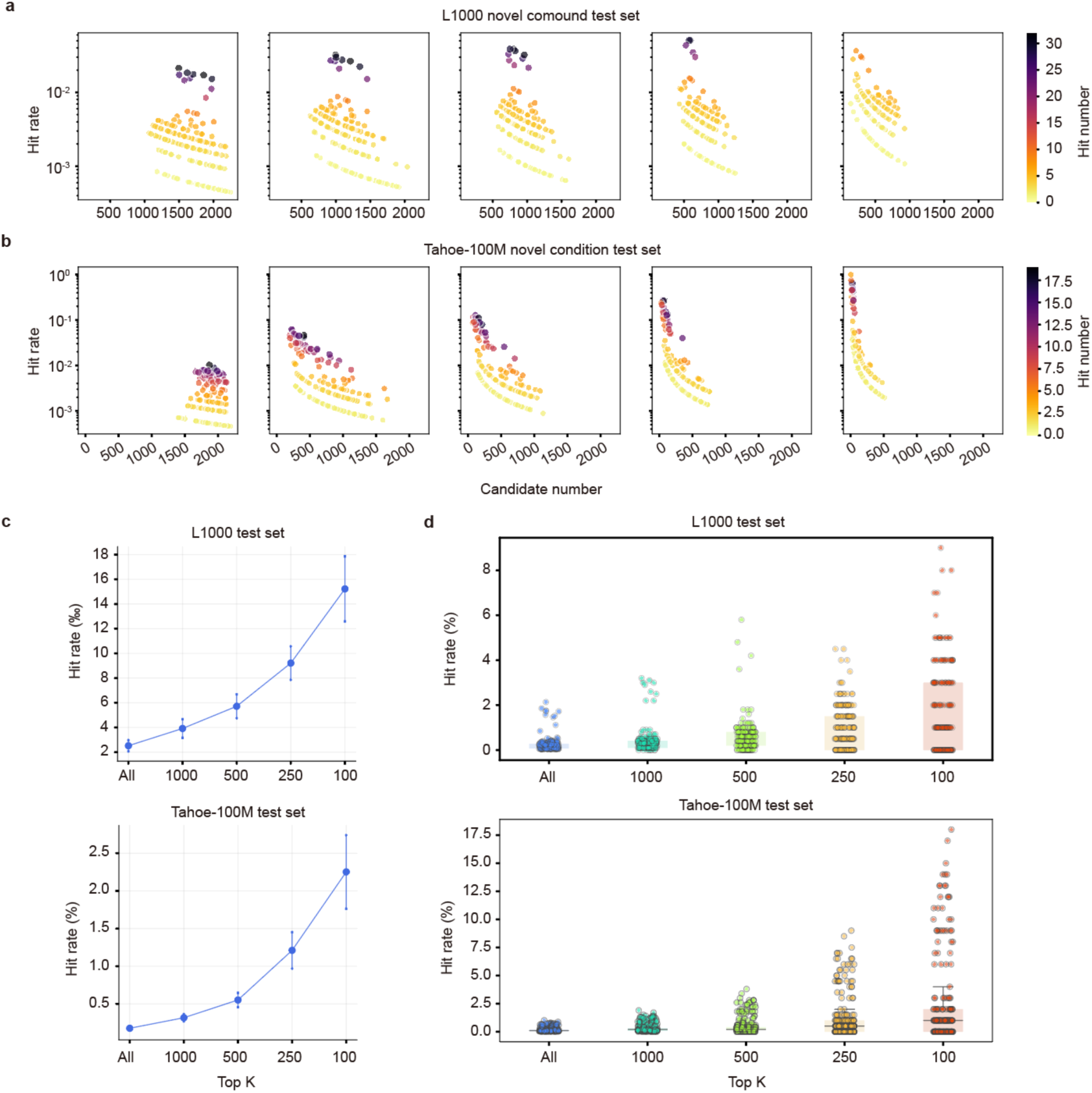
Scoring model prioritizes effective candidates. **a-b**, Effect of filtering by score threshold on candidate retention, hit rate, and hit number in the L1000 (a) and Tahoe-100M (b) test sets. The color is proportional to hit number, with darker indicates more hit number. Y-axis is log-scaled for better representation. **C**, Effect of top-K filtering by score ranking on hit rate in the L1000 (top, N=203) and Tahoe-100M (bottom, N=242) test sets. Error bar, 95% confidence intervals. **d**, Distribution of hit rates per input phenotype filtering by top-K candidates. Centre line, median; box limits, upper and lower quartiles; whiskers, 1.5× interquartile range.

**Figure S6.**
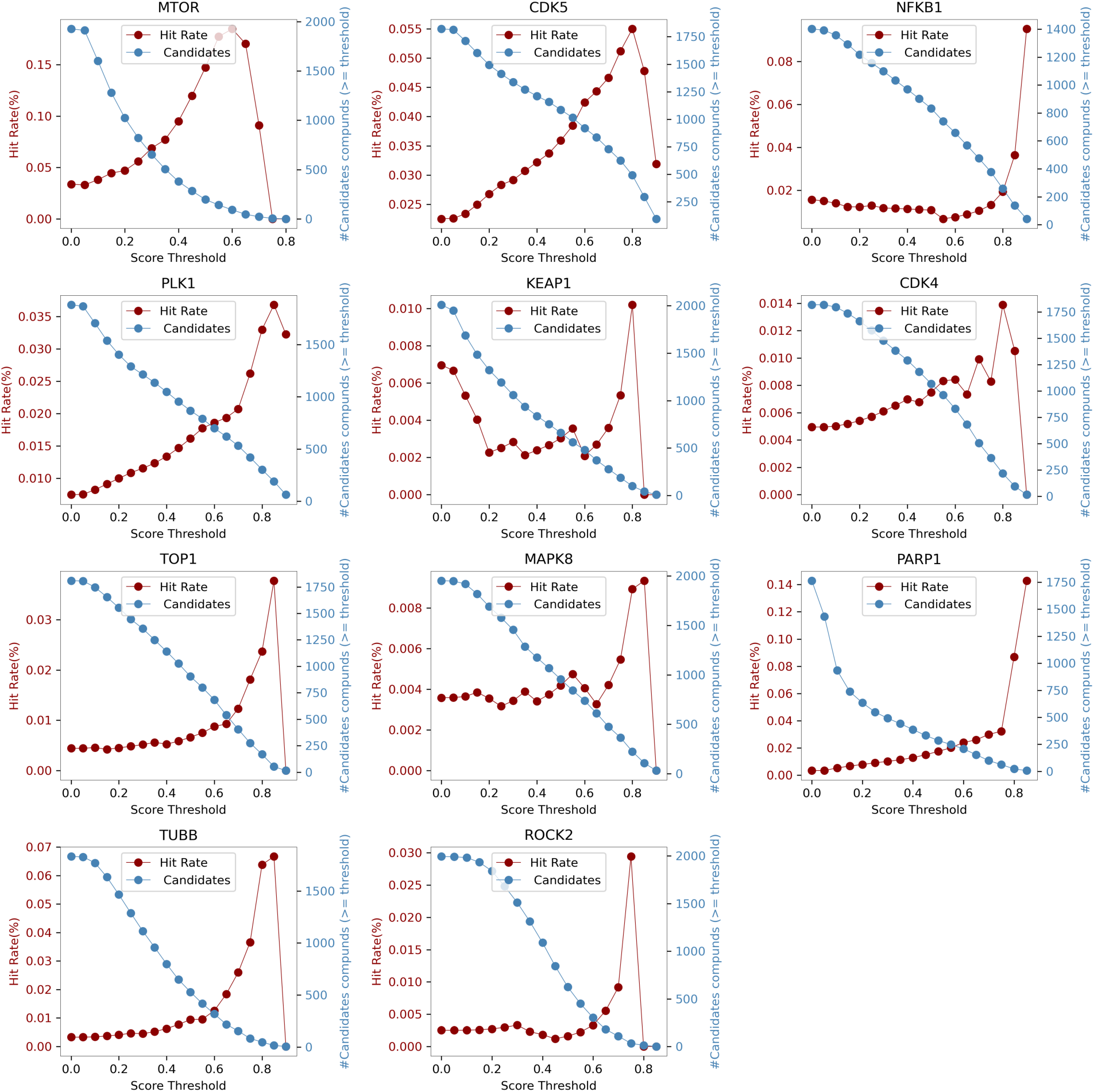
Effect of filtering by score threshold on candidate retention and hit rate in the genetic perturbation test sets.

**Figure S7.**
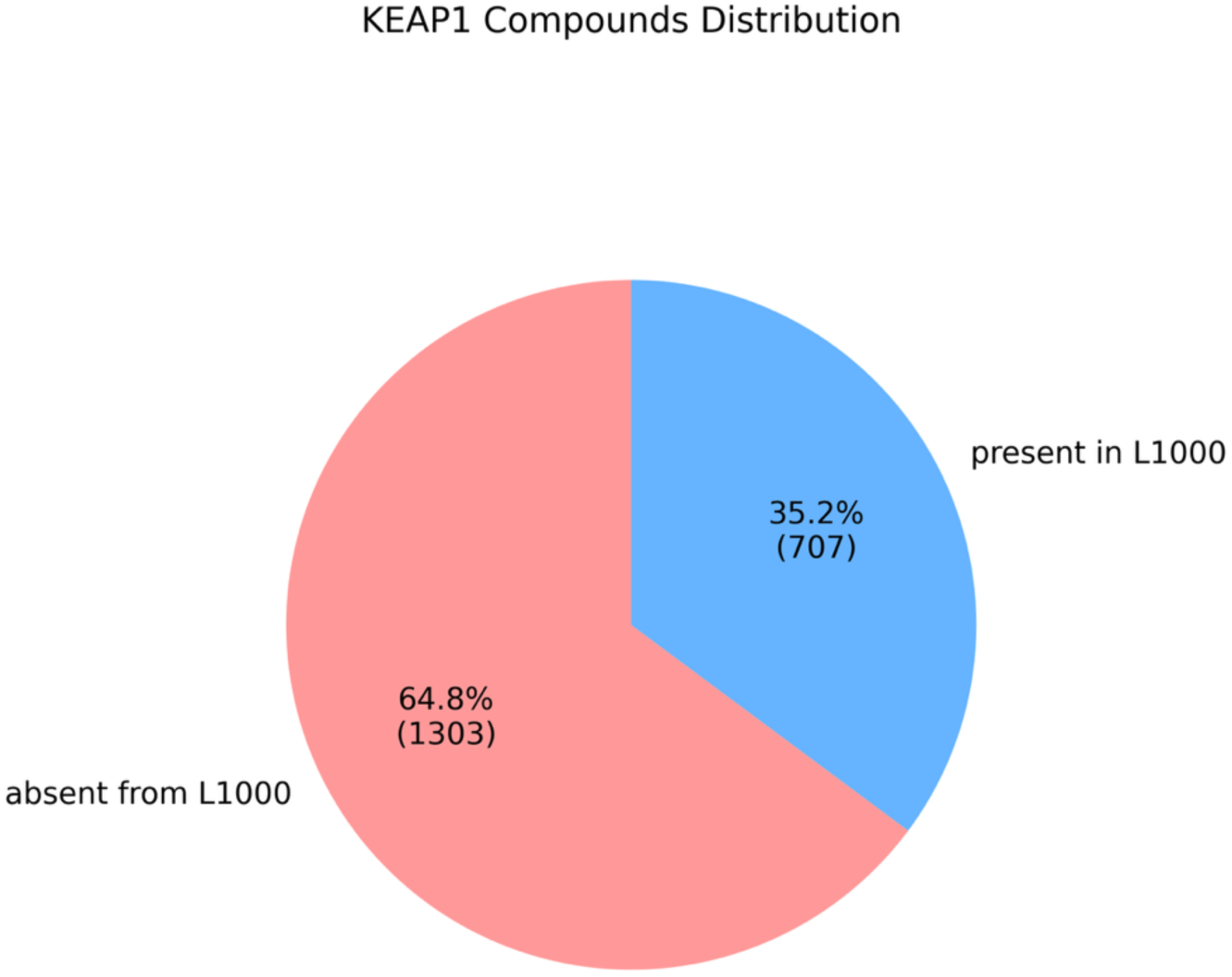
Pie chart showing the proportion of GEMGen-generated compounds that are present in or absent from the L1000 library for the KEAP1 knockdown signature.

**Figure S8.**
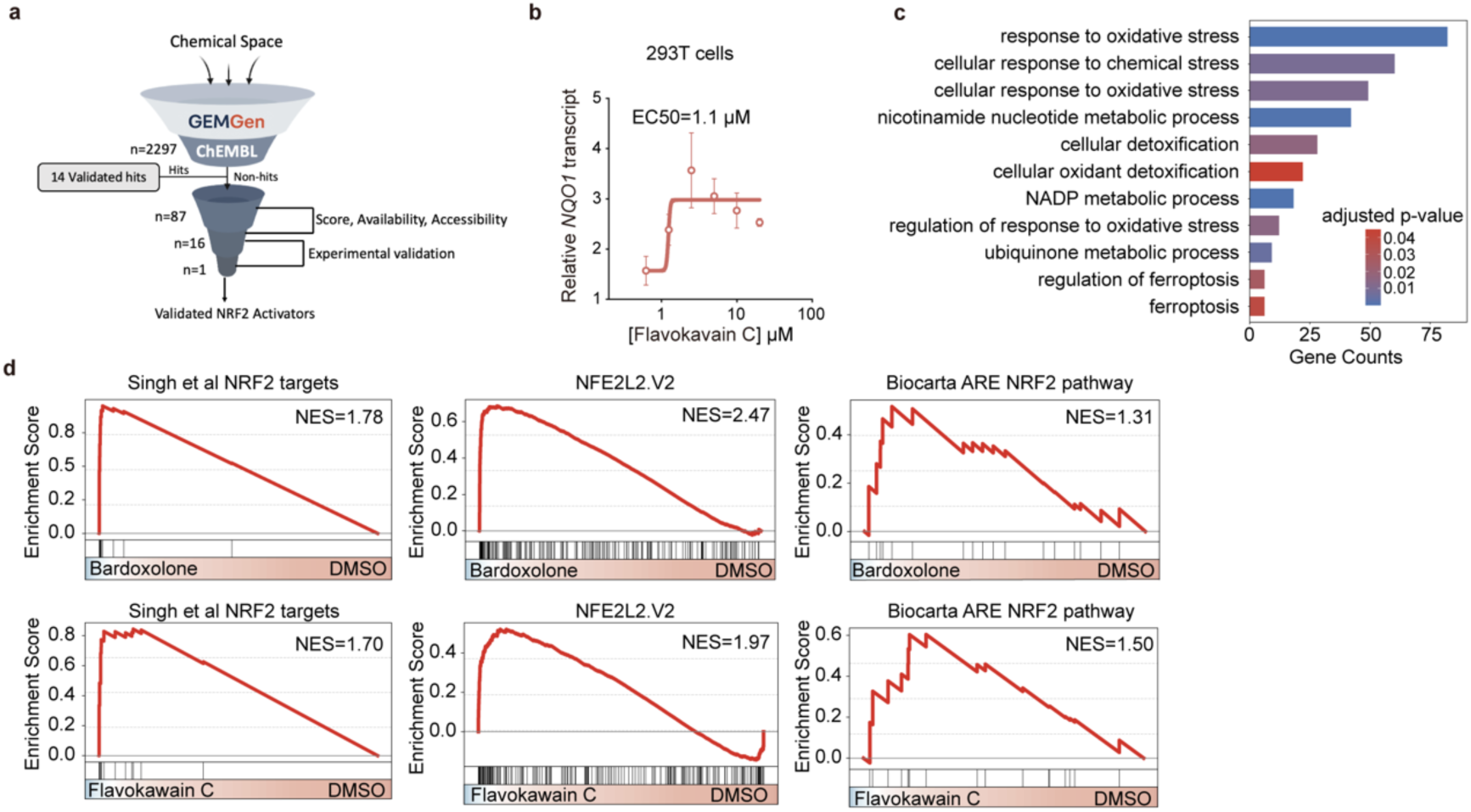
GEMGen generates KEAP1 inhibitor to activate NRF2 signaling. **a**, Schematic illustration of the selection process for experimental validation of KEAP1 inhibitors generated by GEMGen. **b**, qPCR analysis of *NQO1* transcript levels in HEK293T cells treated with different concentrations of flavokawain C. Data are shown as mean ± s.e.m; EC_50_ = 1.1 µM. **c**, Gene ontology analysis of RNA sequencing data from K562 cells treated with 10 µM flavokawain C reveals enrichment in antioxidant response pathways. **d**, Gene-set enrichment analyses of RNA sequencing data from K562 cells treated with flavokawain C or bardoxolone using indicated NRF2 related gene sets. NES, normalized enrichment score.

